# 2’-*O*-methylation alters the RNA secondary structural ensemble

**DOI:** 10.1101/2020.05.28.121996

**Authors:** Hala Abou Assi, Honglue Shi, Bei Liu, Mary C. Clay, Kevin Erharter, Christopher Kreutz, Christopher L. Holley, Hashim M. Al-Hashimi

## Abstract

2’-*O*-methyl (Nm) is a highly abundant post-transcriptional RNA modification that plays important biological roles through mechanisms that are not entirely understood. There is evidence that Nm can alter the biological activities of RNAs by biasing the ribose sugar pucker equilibrium toward the C3’-*endo* conformation formed in canonical duplexes. However, little is known about how Nm might more broadly alter the dynamic ensembles of non-canonical RNA motifs. Here, using NMR and the HIV-1 transactivation response (TAR) element as a model system, we show that Nm preferentially stabilizes alternative secondary structures in which the Nm-modified nucleotides are paired, increasing both the abundance and lifetime of a low-populated short-lived excited state by up to 10-fold. The extent of stabilization increased with number of Nm modifications and was also dependent on Mg^2+^. Through phi (Φ) value analysis, the Nm modification also provided rare insights into the structure of the transition state for conformational exchange. Our results suggest that Nm could alter the biological activities of Nm-modified RNAs by modulating their secondary structural ensembles as well as establish the utility of Nm as a tool for the discovery and characterization of RNA excited state conformations.

## INTRODUCTION

Nm is a highly abundant post-transcriptional modification (Figure 1A) found in non-coding (1-3) and coding RNAs (4,5). It can be added to the ribose sugar moiety of all four nucleotides (Nm = Am, Um, Cm and Gm) (Figure 1A) either *via* stand-alone methyltransferases (6) or by the enzyme fibrillarin when guided by box C/D small nucleolar RNAs (snoRNAs) (7,8). Nm modifications are important for the cellular activity of diverse RNAs (9,10) and have been linked to diseases (11-14). For example, ribosomal RNAs (rRNA) contain more than 100 Nm sites (15), many in the decoding and peptidyl-transferase centers (16). Downregulation of fibrillarin and loss of rRNA Nm sites results in impaired ribosomes that are incapable of translating mRNA (17). Conversely, overexpression of fibrillarin and the concomitant increase in Nm modifications is associated with an increase in protein translation in rapidly dividing breast cancer cells (12). In the spliceosome, all small nuclear RNAs (snRNAs) harbor Nm modifications, some of which are required for proper spliceosome assembly and function (18,19). The loss of Nm modifications on snRNA U6 due to knockout of the La-related protein, which helps load U6 snRNA onto the box C/D snoRNA protein complex (20), changes splicing fidelity, impairs spermatogenesis in mice (20), and contributes to the Alazami syndrome (21). Loss of snoRNA-guided Nm modifications on snRNAs also leads to profound defects in cardiac mRNA splicing and development (13,22,23).

**Figure 1.**
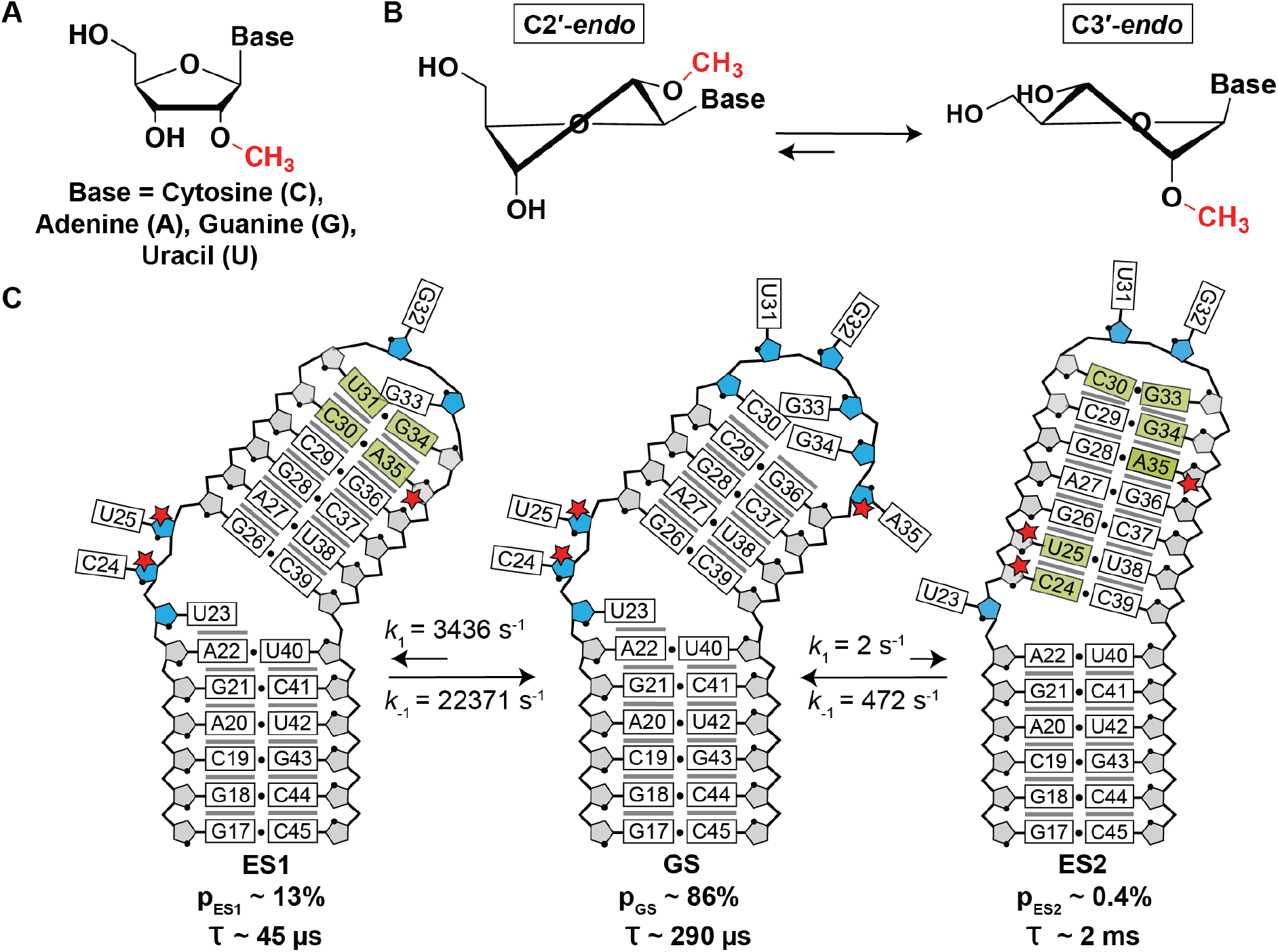
A) Chemical structure of a 2’-*O*-methylated nucleoside (Nm). B) Nm biases the sugar pucker towards C3’-*endo* (34). C) Secondary structure of the TAR GS and ESs. The exchange parameters shown are for TAR in the absence of Mg^2+^ as reported previously (66,67). Nucleotides that undergo sugar repuckering from C2’-*endo* (blue) in the GS to C3’-*endo* (grey) in the ES are indicated using green bases in the ESs. Red stars indicate the Nm-modified residues. Black dots denote base pairing and grey lines denote stacking. The lifetime (τ) of each ES is equal to 1/*k*_-1_, and τ of the GS = 1/(*k*_1,ES1_ + *k*_1,ES2_).

In contrast to the well-studied post-transcriptional modification *N*^6^-methyladenosine (m^6^A) (24-28), which mainly exerts its biological activity by recruiting reader proteins (28-31), no reader proteins have been discovered for Nm to date. Rather, it appears that in some cases, the Nm modification serves to alter the conformational preferences of the Nm-modified RNAs. In isolated mononucleotides and unpaired pyrimidine nucleotides, the ribose moiety exists in a 60:40 dynamic equilibrium between C3’-*endo* and C2’-*endo* sugar pucker conformations (32), whereas most paired nucleotides or nucleotides that stack intra-helically predominantly adopt the C3’-*endo* conformation (33). In pyrimidine nucleosides (Um and Cm) and dinucleotides (UmpU and CmpC), 2’-*O*-methylation biases the sugar pucker equilibrium in favor of C3’-*endo* by 0.1–0.6 kcal/mol (Figure 1B) (34-37). This bias arises due to intra-residue steric repulsion between the 2’-*O*-methyl, the 3’-phosphate, and the 2-carbonyl groups when in the C2’-*endo* conformation (34). The Nm modification has also been shown to stabilize RNA helices by an average of ~0.2 kcal/mol per Nm (38). This is due to a reduction in entropy loss upon duplex formation, as a result of pre-organizing the C3’-*endo* sugar pucker in the single-strand (34,39) and/or favorable enthalpic interactions (40). Since Nm affords increased thermal stability of duplexes and greater stability toward nuclease digestion, it has important applications in FDA-approved oligonucleotide therapeutics (41-44), siRNAs (45,46), and anti-miRNAs (47).

These biases in RNA conformation induced by Nm can serve important biological roles. For example, the bias in sugar pucker introduced by Um at the first anticodon position in tRNA is proposed to reduce misrecognition of noncognate codons, which require a C2’-*endo* conformation (34). In the A loop of the 23S rRNA, the universally conserved Um2552 intercalates between two bases to help maintain the active conformation for the G2553 base, which is directly involved in accommodation of the aminoacyl-tRNA (48). These and other modifications appear to play the role of maintaining distorted RNA conformations at the functional interfaces of the ribosome (48). U2-U6 and U4-U6 snRNAs are critical structural elements of the spliceosome that are rich in Nm modifications (3). These snRNAs undergo changes in secondary structure and base-pairing which are essential for the assembly and disassembly of the spliceosome as well as in cycling between different conformational states required for catalysis (49,50). Nm modifications in the active core of the spliceosome U2-U6 snRNA complex have been shown to promote a conformational transition between a three-way and four-way junction proposed to aid correct positioning of pre-mRNA for exon ligation (51). The modification can also sterically hinder stabilizing interactions between Nm modified mRNAs and ribosomal-monitoring bases, thus disrupting tRNA selection and proofreading and inhibiting translation (5,52).

For the vast majority of the cases however, the role of Nm modifications has remained elusive. While it is known that Nm modifications can stabilize canonical duplexes, little is known about how the modification might more broadly impact the propensities of RNAs to form alternative secondary structures in more complex motifs (such as U2-U6) in which they are frequently found (10,53,54). By biasing sugar pucker and favoring helical conformations, we hypothesized that Nm could preferentially stabilize alternative secondary structures in which the Nm-modified residues are helical with predominantly C3’-*endo* sugar pucker, relative to conformations in which the Nm-modified residues are non-helical and enriched in the C2’-*endo* sugar. These alternative secondary structures, often referred to as ‘excited states’ (ESs) (55-62), are typically transient and low-abundance, but they have been shown to play important roles in the biological activities of RNAs (58,63,64). The ESs of RNAs form by reshuffling base pairs in and around bulges and they tend to be enriched in helical residues, as unpaired nucleotides in bulges and internal loops from the dominant ground state (GS) form mismatches in the ES (Figure 1C). We also hypothesized that the degree to which an ES is stabilized relative to the GS could be controlled by changing the number of nucleotides that are modified. In this regard, it is interesting to note that many RNAs possess clusters of Nm-modified nucleotides (2,53) that could have a synergistic effect on fine-tuning the conformational exchange.

By changing the abundance and possibly the lifetime of ESs, Nm modifications could change RNA ensembles without changing sequence, providing an additional chemical layer with which to optimize RNA structural dynamics for folding and function. In addition to potentially explaining some of the biological roles of Nm, having an ability to control RNA conformational ensembles through Nm modifications may provide a valuable chemical tool for elucidating the characteristics of these low-abundance and short-lived conformations that are difficult to study (55). Here, we have tested our hypothesis using the well-characterized HIV-1 transactivation response element (TAR) RNA as a model system for non-canonical bulge motifs (65). Although TAR is not naturally modified with Nm, its secondary structural ensemble in its unmodified form has been extensively characterized previously using NMR relaxation dispersion (RD) (66,67). It therefore provides an excellent model system for testing our Nm-directed preferential stabilization of paired ESs hypothesis.

Prior NMR RD studies (66,67) showed that TAR exists as a dynamic ensemble involving a GS and at least two ESs (ES1 and ES2) (Figure 1C) (68). ES1 forms rapidly (*k*_ex_ = *k*_1_ + *k*_-1_ ~ 25800 s^-1^) by zipping up the apical loop to form two mismatches, and has a relatively high population (p_ES1_) of ~13% (66). On the other hand, ES2 forms more slowly (*k*_ex_ = 474 s^-1^) through more extensive remodeling of the trinucleotide bulge, upper stem, and apical loop, and has a substantially lower population (p_ES2_ ~0.4%) (67). Based on a recent NMR study (33), the unpaired bulge and apical loop residues C24, U25, and A35 are highly enriched in the C2’-*endo* conformation in the GS whereas they are predominantly C3’-*endo* and helical in ES1 (A35) and ES2 (C24, U25, and A35). We reasoned that introducing Nm at these three nucleotides would stabilize both ESs relative to the GS. Additionally, we hypothesized that the extent of stabilization would be greater for ES2 than ES1, given that all three modifications favor ES2 whereas only Am35 favors ES1.

Indeed, NMR RD measurements showed that Nm increased both the abundance and the lifetime of TAR ES1 and ES2. Through a phi (Φ) value analysis (69-73), the impact of Nm on the exchange kinetics also allowed us to obtain rare conformational insights into the transition state (TS) for the GS-ES conformational exchange. Our results reveal a new mechanism by which Nm could alter the biological activity of Nm-modified RNAs as well as establish the utility of Nm as a chemical tool for the in-depth characterization of alternative ES RNA conformations.

## MATERIAL AND METHODS

### Preparation of RNA samples

Unlabeled TAR, TAR-A35 and TAR-C24U25A35 and ^15^N and ^13^C site-labeled TAR and TAR-C24U25A35 oligonucleotides were synthesized using a MerMade 6 Oligo Synthesizer (BioAutomation) for solid-phase synthesis using standard phosphoramidite chemistry and deprotection protocols (74,75). Unlabeled phosphoramidites were purchased from ChemGenes and site-labeled phosphoramidites were synthesized as described previously (57,76). All RNA samples were prepared using 2’-TBDMS protected phosphoramidites using 1 μmol standard synthesis columns (1000 Å). The final 5’-DMT (4,4’-dimethoxytrityl) was removed during the synthesis for DMT-off deprotection and PAGE purification. Removal of nucleobase protecting groups and cleavage from the 1 μmol columns was achieved using 1 ml of 30% ammonium hydroxide and 30% methylamine (1:1) followed by 2 hours incubation at room temperature. The solution was then air-dried. In order to remove the 2’-TBDMS protecting groups, 100 μL DMSO was added and the samples were heated at 65 °C for 5 min to ensure the samples were fully dissolved. Then 125 μL TEA-3HF was added and the samples were heated at 65 °C for 2.5 h. Samples were then precipitated overnight using 3 M sodium acetate and 100% ethanol, air dried, then dissolved in water for gel purification. All RNA samples were purified using a 20% (w/v) polyacrylamide gel with 8 M urea and 1 X Tris/borate/EDTA. RNA was removed from the excised gel by electro-elution in 1x Tris/acetic acid/EDTA followed by ethanol precipitation. The purified RNA was then annealed in water at a concentration of 50 μM by heating at 95 °C for 5 min followed by cooling on ice for 60 min. The samples were then buffer exchanged five times using an Amicon Ultra-15 centrifugal filter (EMD Milipore) with a 3 kDa cutoff into NMR buffer (15mM sodium phosphate, 25 mM NaCl, 0.1mM EDTA, 10% D2O) at pH 6.4 with or without 1/3 mM Mg^2+^ for the site-labeled/unlabeled samples. The final concentration of unlabeled and site-labeled RNA NMR samples ranged between 0.7-2.0 mM and 0.3-0.7 mM, respectively.

### UV optical melting experiments

UV optical melting experiments were conducted on a PerkinElmer Lambda 25 UV/VIS spectrometer with RTP 6 Peltier Temperature Programmer and a PCB 1500 Water Peltier System. Samples with ~3 μM RNA were prepared by buffer exchanging the RNA into desired buffers (15mM sodium phosphate, 25mM NaCl, 0.1mM EDTA, pH 6.4 with or without 1/3mM Mg^2+^) five times with a centrifugal concentrator (EMD Millipore). Optical melting experiments were performed in triplicates using a sample volume of 400 μL in a Teflon-stoppered 1 cm path length quartz cell. The absorbance at 260 nm (A_260_) was measured as a function of varying the temperature between 15 and 95 °C at a ramp of 1 °C/min. The absorbance curves were fit to two-state model to obtain the thermodynamic parameters using an inhouse Mathematica script and the following equations:

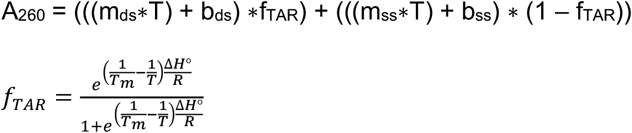

where m_ds_, b_ds_, m_ss_ and b_ss_ are coefficients representing the temperature dependence of the extinction coefficients of the folded and single-stranded species, T is the temperature (K), R is the gas constant (kcal/mol/K), f_TAR_ is the fraction of folded TAR at a given temperature and ΔH^o^ is the enthalpy of annealing. The free energy and entropy of annealing, ΔG^o^ and ΔS^o^ were calculated using the following equation:

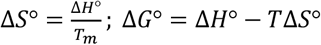

The uncertainty in *T*_m_, ΔH^o^, ΔG^o^, and ΔS^o^ was obtained based on the standard deviation in triplicate measurements. Differences in the standard free energy, entropy, and enthalpy were computed as following:

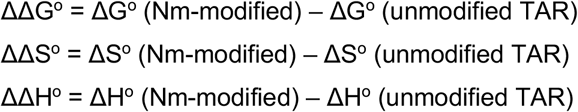

For the triply modified Nm TAR samples with high ES populations, the lower baseline of the absorbance curves was seen to deviate from linearity. Interestingly, a statistically better fit to the data could be obtained using a three-state fit of the absorbance curves while considering the GS, ES and unfolded species as contributing to the measured absorbance (data not shown). Nevertheless, fitting including three-states was seen to minimally influence the stabilities relative to two-states fitting (data not shown). Details of this analysis are outside the scope of this study and will be discussed in future studies.

### NMR Experiments

All NMR experiments were performed on a 600 MHz Bruker NMR spectrometer equipped with an HCN cryogenic probe. Data were processed using NMRpipe (77) and analyzed using SPARKY (T.D. Goddard and D.G. Kneller, SPARKY 3, University of California, San Francisco). Resonances in Nm-modified TAR were assigned based on prior assignments of unmodified TAR (33) and confirmed using 2D HSQC, HMQC, and ^1^H-^1^H NOESY experiments (150 ms mixing time).

#### Determining ES1 population using chemical shift perturbation (CSP) analysis

Because the GS-ES1 exchange is fast on the NMR chemical shift timescale (66), the observed resonance (δ_obs_) in the 2D [^13^C, ^1^H] HSQC spectra of unmodified TAR corresponds to a population-weighted average between the chemical shifts of the GS (δ_GS_) and ES1 (δ_ES1_):

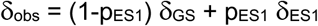

in which 1-p_ES1_ and p_ES1_ are the populations of the GS and ES1, respectively. For unmodified TAR, p_ES1_ ~13% and Δδ = δ_ES1_ - δ_GS_ = 2.6 ppm, as determined previously using ^13^C *R*_1ρ_ RD (66). This allowed determination of δ_GS_ and δ_ES1_ for the various spins that experience RD (C30-C1’, U31-C1’, U31-C6, G34-C1’ and G34-C8). Note that A35-C1’ and A35-C8 are expected to exhibit CSPs towards ES1 chemical shifts; however, these resonances were excluded from analysis since they exhibit sizeable CSPs due to the chemical modification itself as verified using density functional theory (DFT) calculations (Table S1). Based on the δ_obs_ measured for Nm-modified TAR, and the δ_GS_ and δ_ES1_ values deduced for unmodified TAR, p_ES1_ was computed for Nm-modified TAR for each of C30-C1’, U31-C1’, U31-C6, G34-C1’ and G34-C8 (Table S2). Similar p_ES1_ values (average p_ES1_ = 28.7 ± 1.5% in TAR-A35 and 28.8 ± 3.1% in TAR-C24U25A35) were obtained for these spins in the Nm-modified TAR consistent with a concerted shift in population toward ES1. Here the uncertainty represents the standard deviation across the different spins. The same approach was used to estimate p_ES1_ for site-labeled G34-C8 TAR (p_ES1_ = 7.3%) and TAR-C24U25A35 (p_ES1_ = 14.5%) in 1 mM Mg^2+^ using the δ_GS_ and δ_ES1_ values calculated for TAR in the absence of Mg^2+^ (Table S2).

#### ^13^C/^15^N R_1ρ_ experiments

^13^C/^15^N *R*_1ρ_ RD measurements were performed on site-labeled TAR and TAR-C24U25A35 RNA in the presence or absence of 1 mM Mg^2+^ as described previously (55). RD measurements on the unmodified chemically synthesized site-labeled TAR were used to confirm reproducibility with previously published data on uniformly ^13^C/^15^N labeled TAR prepared using *in vitro* transcription (66,67). The spin lock power and offset combinations used in *R*_1ρ_ experiments are listed in Table S7.

#### Analysis of ^13^C/^15^N R_1ρ_ data

The peak intensity in ^13^C/^15^N *R*_1ρ_ measurements at each relaxation delay was extracted using NMRPipe (77). The *R*_1ρ_ value was obtained by fitting the peak intensities at all delay points to a mono-exponential decay and the error was estimated by Monte-Carlo simulation with 500 iterations as described previously (55). The off-resonance *R*_1ρ_ data was fit to a two-state exchange model using the Bloch-McConnel equations and an in-house Python script with the fitting errors estimated by a Monte-Carlo scheme (55). The fitted exchange parameters included the population (p_B_), exchange rate (*k*_ex_), forward rate constant (*k*_1_), backward rate constant (*k*_-1_), and the difference in chemical shift between the ES and GS (Δω = ω_ES_ – ω_GS_). In all cases, the fitting assumed *R*_1,GS_ = *R*_1,ES_ = *R*_1_ and *R*_2,GS_ = *R*_2,ES_ = *R*_2_. In the *R*_1ρ_ plots in Figures 3 and 4, Ω_OBS_=ω_OBS_-ω_RF_, where ω_OBS_ is the Larmor frequency of the observed resonance and ω_RF_ is angular frequency of the applied spin-lock power.

#### GS-ES1 exchange

For the fast exchanging ES1, both in absence and presence of Mg^2+^, the initial alignment of the magnetization was assumed to be along the average effective field of the GS and ES (ω_eff,AVG_) when fitting data to the Bloch-McConnel equations (55). Because the GS-ES1 exchange is fast on the NMR chemical shift timescale (66), and only a single RD probe (G34-C8) was utilized, the exchange parameters, particularly p_ES1_, have high uncertainty (Table S3). Nevertheless, fitting the *R*_1ρ_ data measured for unmodified TAR in the absence of Mg^2+^ assuming the previously reported p_ES1_ (*~* 13%) (66) yielded exchange parameters (*k*_ex_, *k*_1_, *k*_-1_, and Δω) that are within error to those reported previously when using many RD probes and fully labeled TAR (Table S3). When fitting *R*_1ρ_ data for TAR in the presence of 1 mM Mg^2+^, the population was fixed to the value determined by CSP analysis (*i.e*. 7%, Table S2). Because Nm slows down the GS-ES1 exchange, the exchange parameters for TAR-C24U25A35 have lower uncertainty (Table S3). Based on the two-state fit of the *R*_1ρ_ data, Δω_G34-C8_ = 2.4 ± 0.2 ppm for TAR-C24U25A35 in excellent agreement with values measured in unmodified TAR (2.6 ± 0.2 ppm). The similar Δω values indicates that the Nm does not lead to a new ES but rather changes the properties of the pre-existing ES1 (Table S3). The fit revealed that Nm increases the ES1 population ~2-fold, from 13% to 30% in the absence of Mg^2+^ and from 7% to 14% in the presence of Mg^2+^, in excellent agreement with the CSPs (Figure 3B and Table S3).

#### GS-ES2 exchange

Because GS-ES2 is in slow exchange, the initial magnetization was assumed to be aligned along the GS when fitting *R*_1ρ_ data to the Bloch-McConnel equations. Global fitting was carried by sharing p_ES2_ and *k*_ex_ for U23-C6 and U38-N3. Due to degeneracy arising from slow exchange (78), *R*_1ρ_ data measured for TAR in the absence of Mg^2+^ and for TAR-C24U25A35 in the absence and presence of Mg^2+^ was fit by fixing the population to the values determined using CEST (Table S6). Based on a two-state fit of the *R*_1ρ_ data, the Δω values obtained for U23-C6 and U38-N3 were in very good agreement with values measured for unmodified TAR, indicating that the Nm does not lead to a new ES but rather changes the properties of the pre-existing ES2 (Table S6). The addition of 1 mM Mg^2+^ does not significantly affect the chemical shifts of the ES2 probes (U23-C6 and U38-N3) used in this study (Figure S7). Therefore, the weak profiles measured for U23-C6 in the presence of Mg^2+^ (Figure 4B) can be attributed to a reduced Δω due to a Mg^2+^ induced downfield shift in U23-C6 GS chemical shift (Figure S3D). A two-state fit of the U38-N3 data yielded Δω values that are in very good agreement to those measured for TAR in the absence of Mg^2+^ confirming that Mg^2+^ does not lead to a new ES but rather changes the properties of the pre-existing ES2 (Table S6).

#### ^13^C/^15^N CEST experiment

All the ^13^C/^15^N CEST measurements for probing GS-ES2 exchange were performed on site-specific labeled TAR (without Mg^2+^) and TAR-C24U25A35 RNA (with and without 1 mM Mg^2+^) as described previously (78-80). The spin lock power and offset combinations used in CEST measurements are listed in Table S8. The relaxation delay was 0.2 sec for U23-C6 (TAR, without Mg^2+^), U23-C6 (TAR-C24U25A35, without Mg^2+^), U38-N3 (TAR-C24U25A35, without Mg^2+^) and 0.3 sec for U38-N3 (TAR, without Mg^2+^), U23-C6 (TAR-C24U25A35, with Mg^2+^), and U38-N3 (TAR-C24U25A35, with Mg^2+^).

#### Analysis of ^13^C/^15^N CEST data

For each spin lock power and offset, the peak intensities in the 1D ^13^C/^15^N CEST were extracted using NMRPipe (77). The uncertainty in peak intensity was obtained from the intensity variations in the regions of CEST profiles that do not contain any intensity dips (80). The exchange parameters were obtained by fitting the data to the two-state Bloch-McConnel equations using an in-house Python script with the fitting errors estimated by a Monte-Carlo scheme with 100 iterations (78). In all cases, the observed peak was assumed to be the GS peak, as the system is in slow exchange regime (55). ES2 global fitting was carried out by sharing p_B_ and *k*_ex_ between U23-C6 and U38-N3 and assuming *R*_1,GS_ = *R*_1,ES_ = *R*_1_ and *R*_2,GS_ = *R*_2,ES_ = *R*_2_. In the CEST profiles shown in Figure 4 and Figure S5, Ω = ω_RF_-ω_OBS_ where ω_OBS_ is the Larmor frequency of the observed resonance and ω_RF_ is angular frequency of the applied spin-lock power.

#### Density functional theory calculations

Density functional theory (DFT) calculations to compute chemical shifts were performed using Gaussian 09c (Gaussian Inc.) (81) as previously described (82). The starting structures of mononucleotide dA/rA and dT/rU were generated using the *fiber* module of the 3DNA suite (83). The structure of rA/rU with C2’-*endo* sugar was derived from dA/dT by adding a 2’-hydroxl group on the deoxyribose and removing the 7-methyl group from dT using GaussView (https://gaussian.com/gaussview6). The structure of modified rA/rU was generated by furtherly adding a 2’-*O*-methyl group on the ribose. In all cases, the phosphate backbone was truncated using GaussView, thus leaving only the nucleoside motifs with all heavy atoms fixed. Geometry optimization was performed using the B3LYP functional (84) with two runs using the 3-21G (85) and 6-311+G (2d, p) basis sets (86). ^13^C chemical shifts were then calculated using the GIAO method (87) using the B3LYP/6-311+G (2d, p) basis sets. The calculated chemical shifts were then referenced to isotropic magnetic shielding of TMS.

#### Phi (Φ)-value analysis

Φ-value analysis (70) was carried out by computing Φ (Table S5) using the following equation:

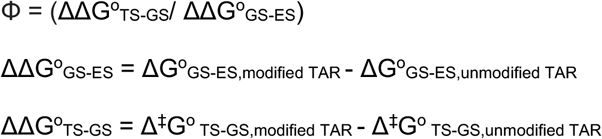

where ΔΔG^o^_GS-ES_ is the change in the free-energy difference between GS and ES induced by the Nm modification obtained from ES population (p_ES_) obtained using *R*_1ρ_ and CEST experiments (Tables S3 and S6):

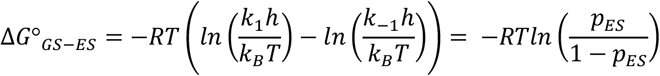

Where R is the gas constant, T is temperature, *h* is Planck’s constant, *k*_B_ is Boltzmann’s constant, *k*_1_ is the forward rate constant, and *k*_-1_ backward rate constant.

ΔΔG^o^_TS-GS_ is the change in the forward free-energy barrier (Δ^‡^G^o^) induced by Nm calculated using the forward rate constant *k*_1_ obtained from *R*_1ρ_ and CEST experiments (Tables S3 and S6):

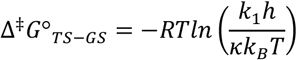

where *κ* is the transmission coefficient and is assumed to be unity.

## RESULTS

### Nm modifications have a small impact on the thermal stability of the TAR GS

We first used UV melting experiments to assess the impact of the modification on the overall thermal stability of TAR, which is dominated by the GS (population ~ 86%) and to a lesser extent ES1 (population ~13%) (see Methods). Given that other post-transcriptional modifications such as m^6^A have been shown to affect RNA thermal stability and structural dynamics in a Mg^2+^-dependent manner (78,88,89), and that the TAR bulge binds Mg^2+^ in a unique conformation (90,91), experiments were performed in the absence and presence of Mg^2+^.

Using solid-phase oligonucleotide synthesis, TAR samples were prepared in which A35 alone (TAR-A35) or in conjunction with C24 and U25 (TAR-C24U25A35) were Nm modified (Figure 1C). A third reference sample was also prepared lacking any modifications (TAR). The single Am35 modification had a negligible effect on the free energy of melting (Table 1 and Figure 2A) both in the absence (ΔΔG^o^ ~ 0.1 kcal/mol) and presence (ΔΔG^o^ ~ +0.2 kcal/mol) of 3mM Mg^2+^. The triple modification slightly stabilized TAR by ΔΔG^o^ ~ −0.3 kcal/mol in the absence of Mg^2+^ and resulted in an equally small degree of destabilization (ΔΔG^o^ ~ +0.4 kcal/mol) in the presence of 3 mM Mg^2+^ (Table 1 and Figure 2A). The addition of 3 mM Mg^2+^ stabilized both Nm-modified and unmodified TAR samples by a more significant amount of ~3 kcal/mol (Table 1) consistent with prior studies (90,91) showing that TAR interacts favorably with divalent metals.

**Table 1.**
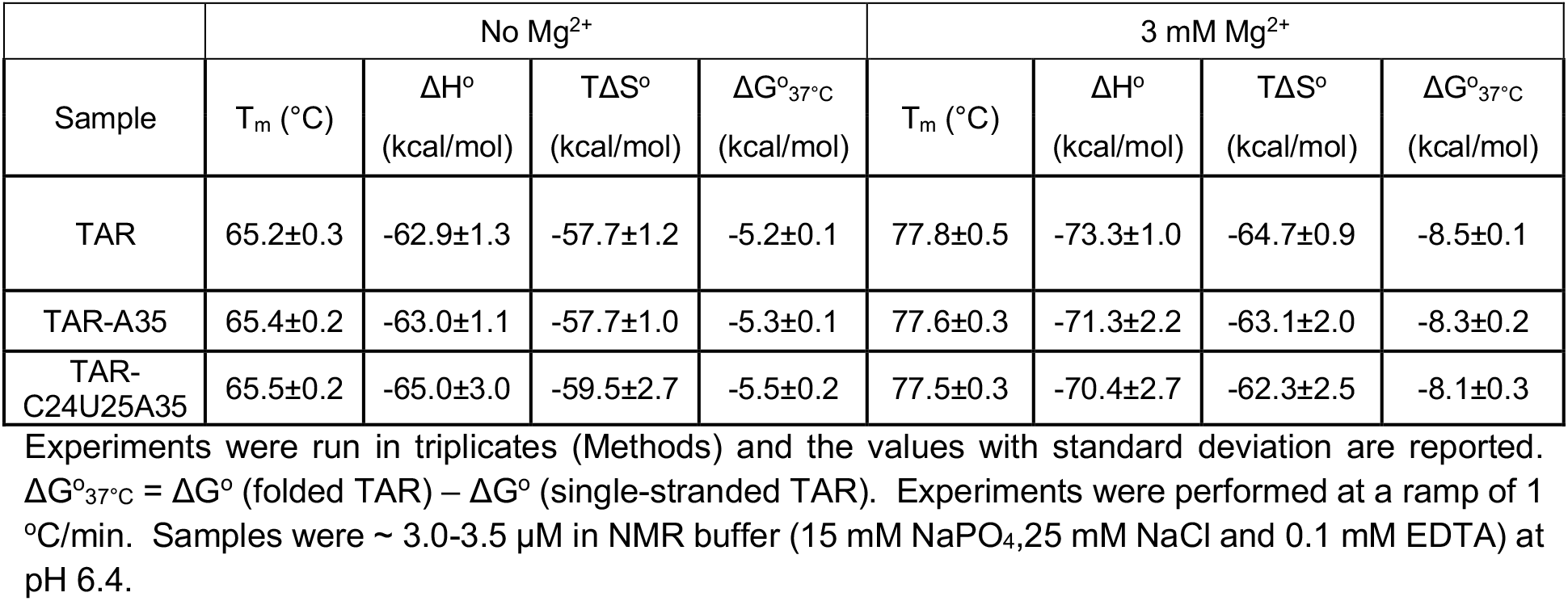
Melting temperature and thermodynamic parameters derived from melting curves in the absence and presence of Mg^2+^.

| | No $Mg^{2+}$ | | | | 3 mM $Mg^{2+}$ | | | |
| --- | --- | --- | --- | --- | --- | --- | --- | --- |
| Sample | $T_m$ (°C) | $\Delta H^\circ$<br>(kcal/mol) | $T\Delta S^\circ$<br>(kcal/mol) | $\Delta G^\circ_{37^\circ C}$<br>(kcal/mol) | $T_m$ (°C) | $\Delta H^\circ$<br>(kcal/mol) | $T\Delta S^\circ$<br>(kcal/mol) | $\Delta G^\circ_{37^\circ C}$<br>(kcal/mol) |
| TAR | 65.2±0.3 | -62.9±1.3 | -57.7±1.2 | -5.2±0.1 | 77.8±0.5 | -73.3±1.0 | -64.7±0.9 | -8.5±0.1 |
| TAR-A35 | 65.4±0.2 | -63.0±1.1 | -57.7±1.0 | -5.3±0.1 | 77.6±0.3 | -71.3±2.2 | -63.1±2.0 | -8.3±0.2 |
| TAR-C24U25A35 | 65.5±0.2 | -65.0±3.0 | -59.5±2.7 | -5.5±0.2 | 77.5±0.3 | -70.4±2.7 | -62.3±2.5 | -8.1±0.3 |

Experiments were run in triplicates (Methods) and the values with standard deviation are reported. $\Delta G^\circ_{37^\circ C} = \Delta G^\circ$ (folded TAR) – $\Delta G^\circ$ (single-stranded TAR). Experiments were performed at a ramp of 1 °C/min. Samples were ~ 3.0-3.5 $\mu M$ in NMR buffer (15 mM $NaPO_4$ , 25 mM NaCl and 0.1 mM EDTA) at pH 6.4.

**Figure 2.**
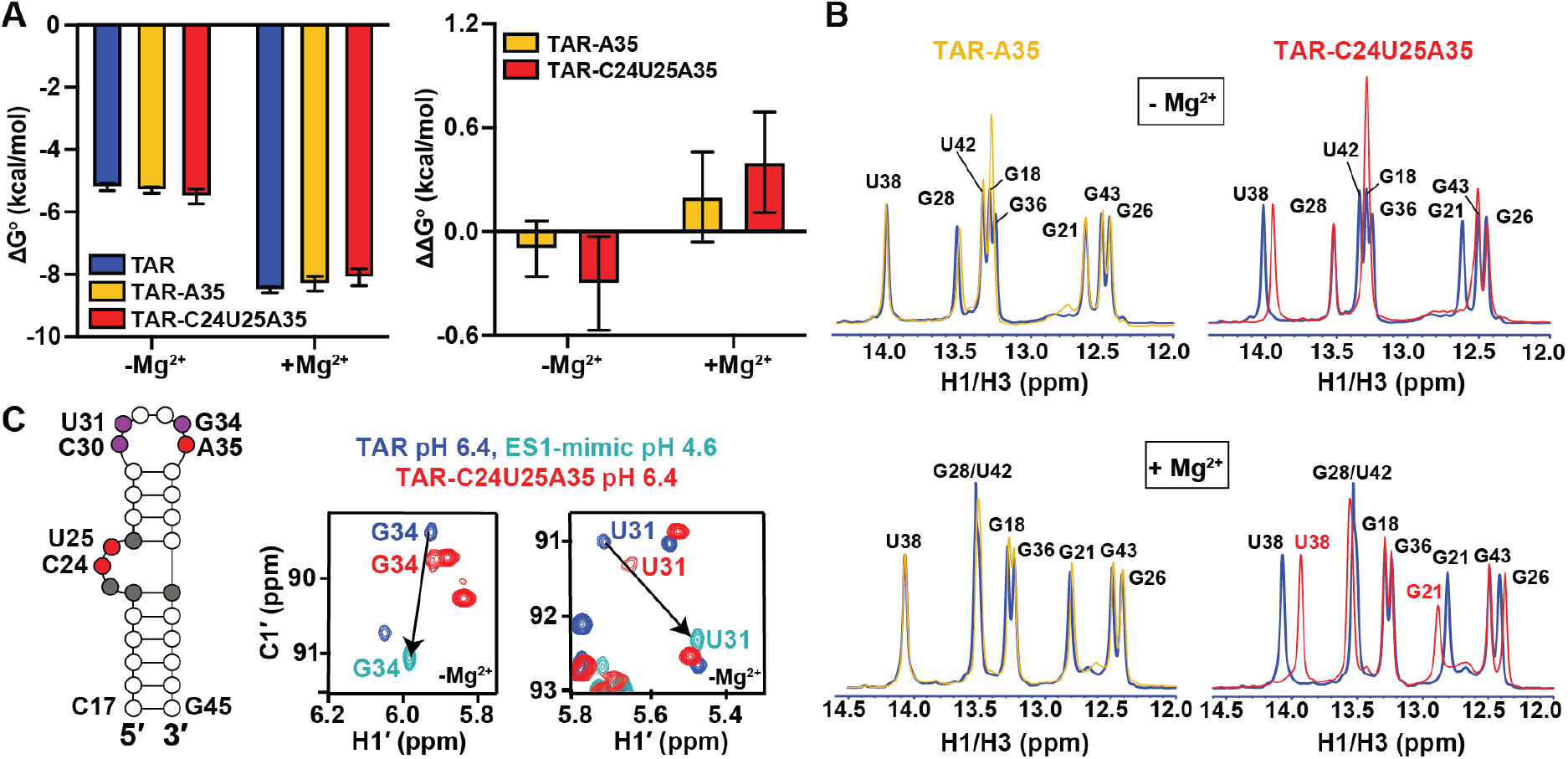
Impact of Nm on the thermal stability of TAR GS. A) ΔG^o^ and ΔΔG^o^ values derived from melting curves in the absence and presence of 3 mM Mg^2+^. Error bars on ΔG^o^ denote standard deviation of triplicate measurements as described in the methods. Error bars on ΔΔG^o^ were obtained by propagating the errors from triplicate measurements (see Methods). B) Overlays comparing 1D ^1^H imino spectra for unmodified TAR (blue), TAR-A35 (yellow) and TAR-C24U25A35 (red) in the absence and presence of 3 mM Mg^2+^. All samples were unlabeled. C) Secondary structure of TAR with Nm-modified residues shown in red, residues showing CSPs towards ES1 are in purple, and residues exhibiting CSPs due to bulge modifications are in grey. Overlay of natural abundance 2D [^13^C, ^1^H] HSQC spectra for the CT-HT region in TAR (pH 6.4), TAR ES1-mimic (pH 4.6) (66), and TAR-C24U25A35 (pH 6.4), in the absence of Mg^2+^. Lowering the pH to 4.6 shifts the population of TAR (structure shown in Figure S2) to stabilize ES1 as the dominant conformation due to protonation of the A35-C30 mismatch unique to ES1 (66). Spectra with complete assignments are shown in Figures S1 and S2.

**Figure 3.**
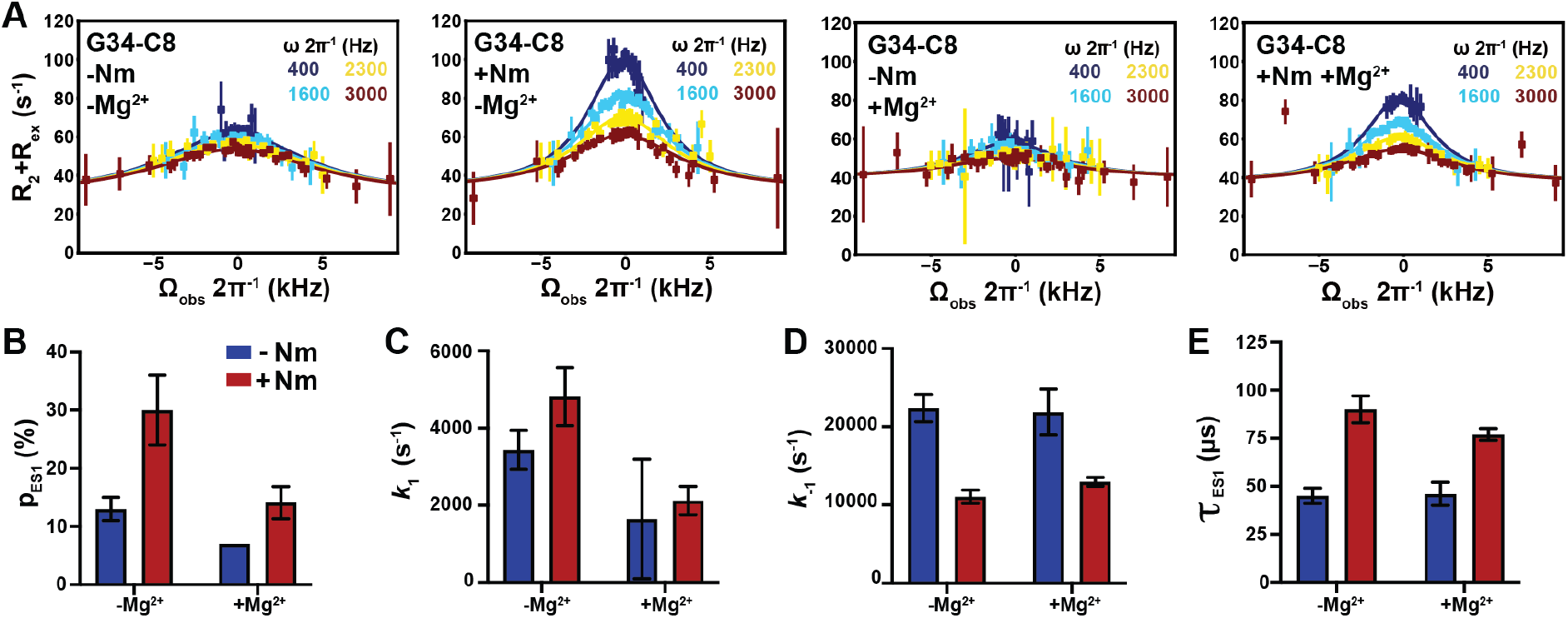
Impact of Nm on GS–ES1 exchange. A) Off-resonance *R*_1ρ_ ^13^C RD profiles for G34-C8 in TAR and TAR-C24U25A35 at 25°C, in the absence and presence of 1 mM Mg^2+^. *R*_1ρ_ data was fit with a two-state model using Bloch-McConnell equations. Spin-lock powers are color-coded. Errors in the *R*_1ρ_ data were calculated as described previously (63). B-E) Comparison of ES1 population (p_ES1_), forward (*k*_1_) and backward (*k*_-1_) rate constants, and lifetime (τ) between TAR and TAR-C24U25A35, in the absence and presence of 1 mM Mg^2+^, as obtained from fitting *R*_1ρ_ data. Error bars were calculated using a Monte-Carlo scheme (63).

These results show that even three Nm modifications have minor contributions to the overall thermal stability of RNAs when placed on unpaired flexible nucleotides. Such small energetic perturbations on the GS make the modification particularly attractive for carrying out Φ-value analysis to gain insights into the transition-state (TS) (69-73). As we show below, while the modifications only weakly affected the energetic stability of the TAR folded state relative to the unfolded one, they had a bigger effect on the energetic stabilities of the two ESs relative to the GS, particularly for ES2.

### Nm modifications impact the TAR conformational preferences

Comparison of the 1D ^1^H NMR spectra of TAR-A35 and TAR-C24U25A35 with their unmodified counterparts (Figure 2B), revealed that Nm minimally impacts the secondary structure of the dominant GS both in the presence and absence of 3 mM Mg^2+^. Both Nm-modified samples showed the imino resonances belonging to the TAR Watson-Crick base-pairs (bps) (Figure 2B). The modifications did however induce chemical shift perturbations (CSPs) in 1D ^1^H NMR (Figure 2B) and 2D [^13^C,^1^H] HSQC spectra (Figure 2C and Figures S1-S3), indicating that they do indeed affect the conformational dynamics of TAR. As described below, the CSPs observed for apical loop residues in the 2D [^13^C,^1^H] HSQC spectrum are as expected if the Nm modifications were to increase the abundance of ES1. Other sizeable CSPs observed at the sugar resonances of the Nm-modified residues (Figures S1-S3) can be attributed to the chemical modification itself, as verified using density functional theory (DFT) calculations (Table S1) (92). Small CSPs were also observed in TAR-C24U25A35 at residues in and around the bulge (grey residues in Figure 2C), indicating a subtle change in the GS conformational ensemble.

### Nm increases the abundance of ES1 relative to the GS

The GS-ES1 exchange is fast on the NMR chemical shift timescale. Consequently, if the Nm modification at A35 were to increase the abundance of ES1 without substantially slowing the exchange kinetics, we would expect to observe CSPs in apical loop residues that are directed specifically towards the ES1 ^1^H and ^13^C chemical shifts, which have been determined previously (66). Furthermore, we would expect similar CSPs for both TAR-A35 and TAR-C24U25A35 given that only Am35 is expected to bias the equilibrium in favor of ES1 (Figure 1C). The magnitude of the shift should be proportional to the degree to which the modification increases the ES1 population (66).

Indeed, for both TAR-A35 and TAR-C24U25A35 in the absence of Mg^2+^, ^1^H and ^13^C CSPs were observed at C30-C1’, U31-C1’, U31-C6, G34-C1’ and G34-C8 that are specifically directed towards the ES1 chemical shifts (Figure 2C and Figure S2). As a negative control, no CSPs were observed at apical loop residues G32 and G33, which have similar chemical shifts in the GS and ES1 as they remain unpaired in the two cases (33) (Figure S2). Based on the magnitude of the CSPs (see Methods), Nm increases the ES1 population ~2-fold from 13 ± 2% to 29 ± 3% in both TAR-A35 and TAR-C24U25A35 (Table S2). The ~0.6 kcal/mol stabilization of ES1 relative to the GS (Figure S4) is larger than the ~0.2 kcal/mol stabilization reported previously from incorporating a single Am into RNA duplexes (93). This is not surprising given that the net effect of Nm on thermodynamic parameters is context-dependent (93). The Am35 modification at the Am35^+^-C30 mismatch could have a more stabilizing effect on ES1 and/or the modification may also destabilize the unpaired conformation of A35 in the GS.

To assess the ES1 abundance in the presence of Mg^2+^, we prepared unmodified TAR and TAR-C24U25A35 samples site-labeled (57,76) with ^13^C at G34-C8 to probe ES1, and with ^13^C at U23-C6 and ^15^N at U38-N3 to probe ES2 (66) (see below). This was necessary to boost NMR sensitivity given that Mg^2+^ significantly broadened the resonances in the 2D NMR spectra (94) (Figure S3A-C). The addition of 1 mM Mg^2+^ to TAR resulted in CSPs at G34-C8H8 that are directed away from ES1 toward the GS, as expected if Mg^2+^ were to stabilize the GS relative to ES1 (Figure S3D). Based on the magnitude of these CSPs, Mg^2+^ reduced the abundance of ES1 two-fold (from 13% to ~7%, Table S2) corresponding to 0.4 kcal/mol destabilization of ES1 relative to the GS (Figure S4). Thus, Mg^2+^ preferentially binds and stabilizes the apical loop in the more open GS conformation relative to the zipped up ES1.

Although Mg^2+^ decreased the ES1 population, introducing Nm once again induced CPSs directed towards ES1 (Figure S3D). Based on these CSPs, Nm increased the abundance of ES1 twofold (from 7% to ~14%, Table S2). Therefore, Nm stabilizes ES1 relative to GS in a manner independent of Mg^2+^, whereas Mg^2+^ stabilizes the GS relative to ES1 in a manner independent of Nm (Figure S3D and Figure S4). This independence could arise because the Mg^2+^ ions that stabilize the GS relative to the ES1 interact with apical loop residues distant from the modified A35 site and are therefore unaffected by the Nm modification.

### Nm increases the lifetime of ES1 in the absence and presence of Mg^2+^

To further confirm that Nm stabilizes ES1, and to gain insights into how Nm affects the GS-ES1 exchange kinetics, we preformed off-resonance spin relaxation in the rotating frame (*R*_1ρ_) RD NMR experiments on the isotopically labeled unmodified TAR and modified TAR-C24U25A35 samples (Figure 3A). The Nm modification and Mg^2+^ had opposite effects on the *R*_1ρ_ profiles measured for G34-C8. Nm enhanced the *R*_1ρ_ profile whereas the addition of Mg^2+^ diminished it (Figure 3A). In accord with the CSP analysis, a two-state fit of the RD data showed that Nm increased the ES1 population ~2-fold both in the presence and absence of Mg^2+^, whereas Mg^2+^ decreased the population by a similar amount in the presence and absence of Nm (Figure 3B and Table S3).

**Figure 4.**
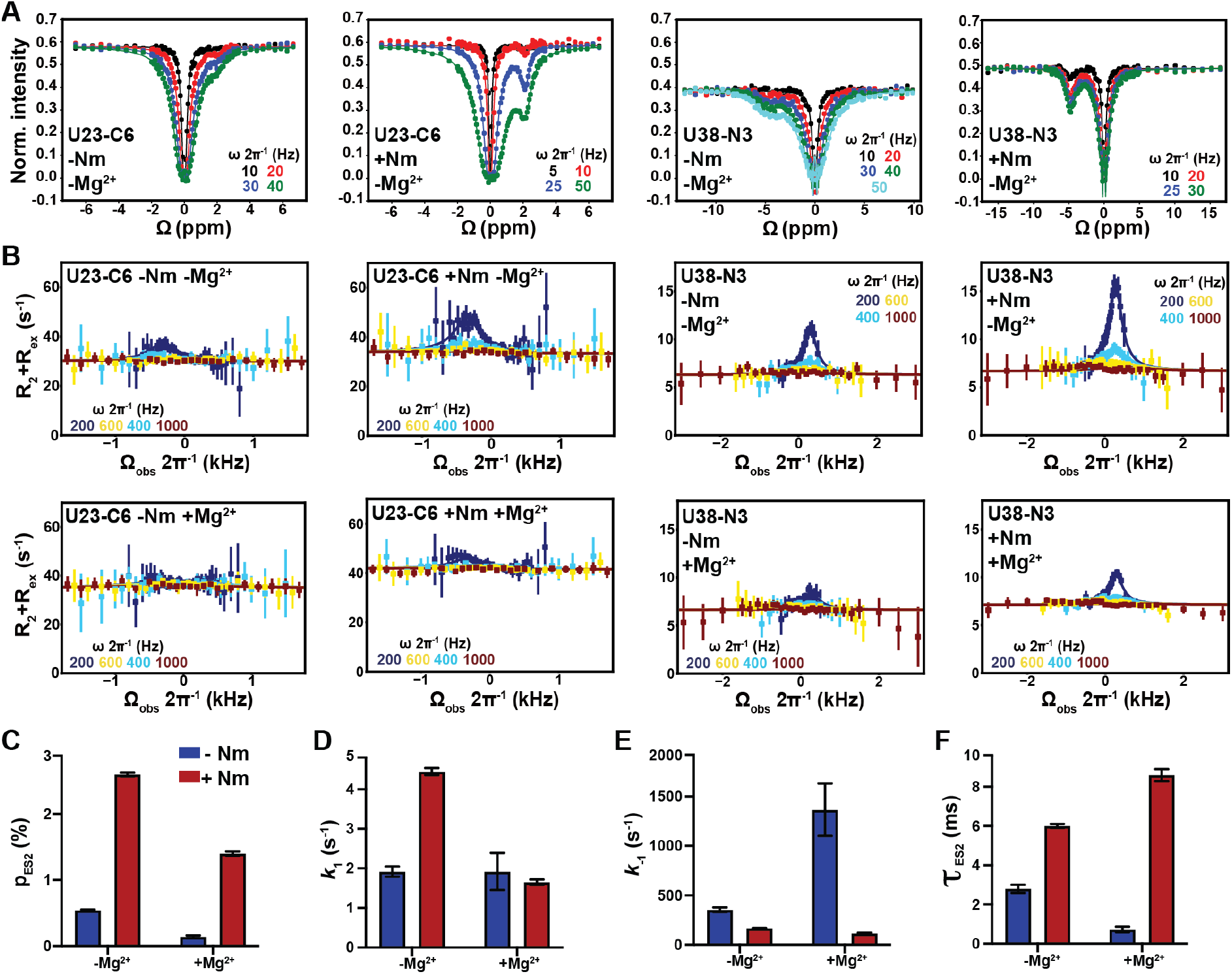
Impact of Nm on GS-ES2 exchange in TAR. A) ^13^C and ^15^N CEST profiles in TAR and TAR-C24U25A35 in the absence of Mg^2+^. B) Off-resonance ^13^C and ^15^N *R*_1ρ_ RD profiles for U23-C6 and U38-N3 in TAR and TAR-C24U25A35 in the absence and presence of 1 mM of Mg^2+^. *R*_1ρ_ data was fit with a two-state model using Bloch-McConnell equations. Fits of *R*_1ρ_ profiles were preformed fixing the population to the value measured using CEST. Errors in the *R*_1ρ_ data were calculated as described previously (63). Spin-lock powers in CEST and *R*_1ρ_ are color-coded. C-F) Comparison of p_ES2_, *k*_1_ and *k*_-1_ rate constants, and τ _ES2_ between TAR and TAR-C24U25A35 in the absence and presence of 1 mM Mg^2+^ obtained from fitting the CEST data. Error bars were calculated using a Monte-Carlo scheme (55).

The RD data also revealed that Nm did not significantly change the forward rate (Figure 3C). Rather, it robustly reduced the backward rate (*k*_-1_) (Figure 3D) and increased the ES1 lifetime by ~2-fold in the absence and presence of Mg^2+^ (Figure 3E). Conversely, Mg^2+^ robustly increased the abundance and lifetime of the GS in the absence and presence of Nm modifications. Thus, the opposing effects of Nm and Mg^2+^ on the GS-ES1 exchange are independent of one another at both the thermodynamic as well as the kinetic level. While this suggests that the modification similarly impacts the energetic stabilities of the GS and the transition state for ES1 (TS1) (Table S4 and Figure S4), the nature of the TS1 was difficult to ascertain given the large uncertainty in the Φ-value (70) (Table S5).

### Nm increases the abundance and lifetime of ES2 in a Mg^2+^-dependent manner

We next turned our attention to how Nm impacts the GS-ES2 exchange using U23-C6 and U38-N3 as RD probes. Here, we expected a greater effect because all three Nm modifications should energetically favor ES2 relative to the GS. To overcome a degeneracy arising due to slow exchange kinetics (Figure S5), we supplemented the *R*_1ρ_ data with measurements of chemical exchange saturation transfer (CEST) experiments (79,80), which are better suited for studying slower GS-ES exchange process (78).

We first benchmarked the unmodified site-labeled TAR sample in the absence of Mg^2+^. As expected, the exchange parameters obtained by fitting CEST (Figure 4A) and *R*_1ρ_ (Figure 4B) data measured for U23-C6 and U38-N3 (Figure 4C-F and Table S6) were in excellent agreement with the values reported previously (67). As predicted, relative to unmodified TAR, TAR-C24U25A35 showed enhanced CEST (Figure 4A) and *R*_1ρ_ profiles (Figure 4B), consistent with an increase in ES2 population (Figure 4C). A two-state fit of the RD data revealed that Nm increased the ES2 population ~5-fold (from ~0.5% to ~2.6%), stabilizing ES2 relative to the GS by ~1 kcal/mol (Table S6 and Figure S6). The ~2.5-fold greater stabilization of ES2 relative to ES1 was as expected from application of three versus a single Nm modification. As with ES1, the Nm modifications increased ES2 lifetime by ~2-fold from ~2.8 to ~6.0 ms (Table S6). However, unlike for ES1, the Nm modifications also increased the forward rate ~2.3-fold (Figure 4D and Table S6). This results in an intermediate Φ-value of ~0.5 for Nm-modified TAR in the absence of Mg^2+^, which is difficult to interpret to obtain structural insights into the transition state for ES2 formation (TS2) (Table S5).

As with ES1, the addition of Mg^2+^ to unmodified TAR diminished the CEST and *R*_1ρ_ profiles of U38-N3 and U23-C6 (Figure 4B) due to a ~4-fold reduction in ES2 population (from ~0.54 to ~0.14%, Table S6). The greater degree to which Mg^2+^ stabilizes the GS relative to ES2 versus ES1 is as expected given that a change in the bulge conformation is required to form ES2 and that four divalent metal ions specifically bind and stabilize the TAR bulge in the GS conformation (91).

When combined together, Nm and Mg^2+^ produced an unexpected result. Based on the *R*_1ρ_ and CEST profiles, introducing the Nm modifications in the presence of Mg^2+^ increased the abundance of ES2 by 10-fold (Figure 4C, 4F and Table S6). The greater stabilization of ES2 relative to the GS in the presence (ΔΔG^o^ = 1.4 kcal/mol) versus absence of Mg^2+^ (ΔΔG^o^ = 1.0 kcal/mol) might in part be explained by the UV melting data (Figure 2A and Table 1), where Nm slightly destabilizes TAR GS in the presence of Mg^2+^ by ~0.4 kcal/mol. Consequently, the Nm modifications at the bulge could potentially be disrupting favorable Mg^2+^ interactions, thus destabilizing the GS and stabilizing the helical conformations formed in the ES2. This underscores the importance of examining how the Nm modifications may impact the conformational properties of non-canonical motifs.

Strikingly, in the presence of Mg^2+^, the introduction of the three Nm sites minimally impacted the forward rate (1.2-fold), whereas they decreased the backward rate by 12-fold and increased the ES2 lifetime by the same amount. Compared to the unmodified TAR ES2 in the presence of Mg^2+^, the addition of Nm leads to a Φ-value of −0.07 ± 0.10 (Table S5) that strongly implies an early transition state for ES2 formation. These results indicate that in the presence of Mg^2+^, the Nm modified bulge residues C24 and U25 remain unpaired and enriched in C2’-*endo* sugar pucker in the TS2 as is observed in the GS.

### Direct observation of ES2 in Nm-modified TAR

If three Nm modifications do indeed increase the population of ES2 to >1%, it should be possible to directly observe ES2 in the NMR spectra of TAR-C24U25A35, given that the GS-ES2 exchange is slow on the NMR chemical shift timescale, and is further slowed down by the Nm modifications. Indeed, a minor imino resonance was observed in 2D [^15^N,^1^H] HSQC spectra of the site-labeled TAR-C24U25A35 but not in unmodified TAR that could be assigned to U38 when forming the wobble U38-U25 mismatch in ES2 both in the absence (Figure 5A) and presence of 1 mM Mg^2+^ (Figure 5B). This resonance shows excellent overlap in both ^1^H and ^15^N chemical shifts with the imino resonance for U38 in a previously described unmodified ES2-mutant (UUCG-ES2) (67) which stabilizes ES2 as the major conformation (Figure S7). Based on integrated volumes of the GS and ES2 resonances, the ES2 population is estimated to be ~3% and ~1% in the absence and presence of Mg^2+^, respectively, in excellent agreement with the ES2 populations (~3% and ~1%, respectively) obtained from the CEST and *R*_1ρ_ data. Thus, Nm increases the abundance of ES2 to a degree that makes it directly observable in NMR 2D spectra.

**Figure 5.**
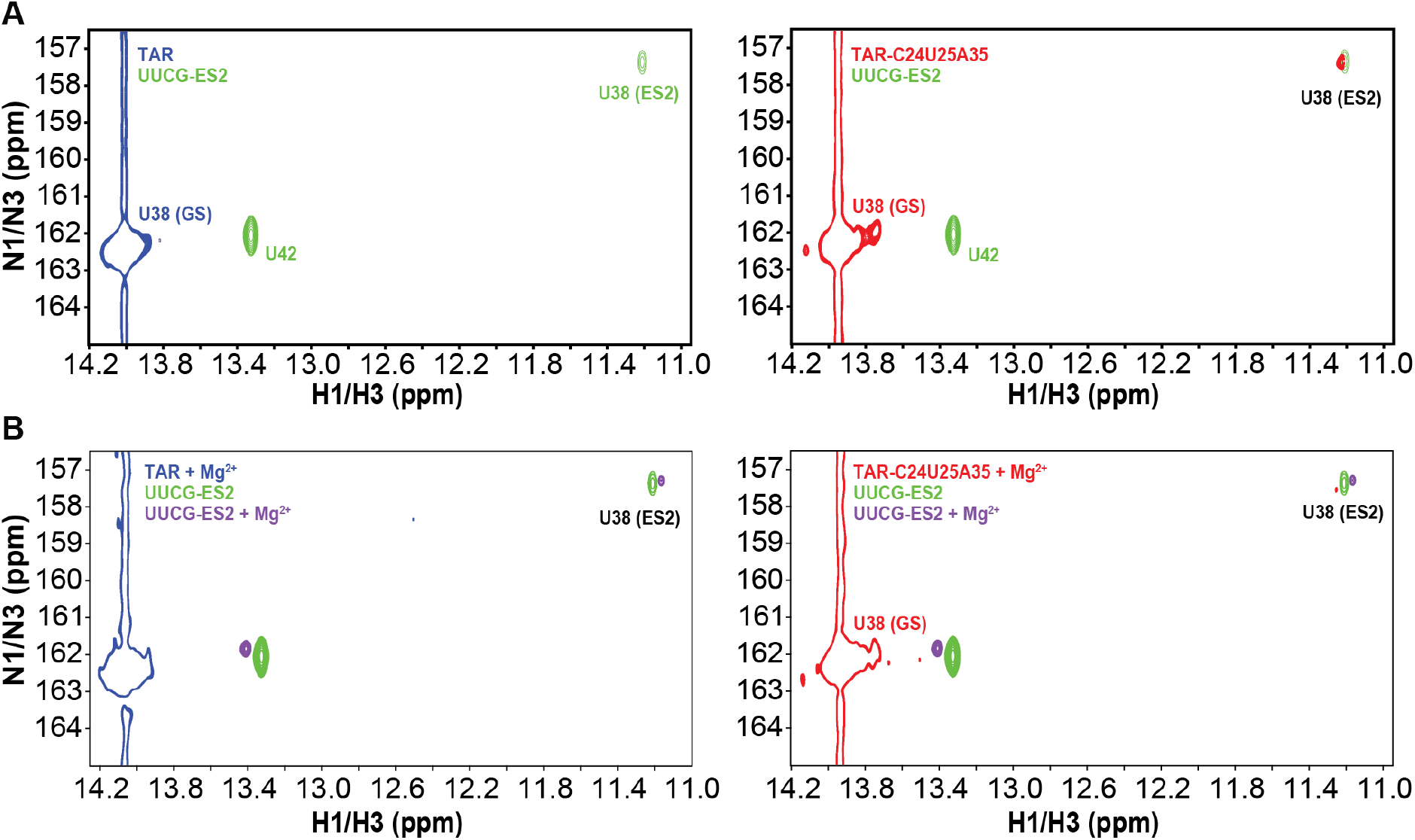
Direct observation of ES2 U38-N3H3 resonance in the Nm-modified TAR sample in NMR 2D spectra. Overlay of 2D [^1^H,^15^N] HSQC spectra of uniformly ^15^N/^13^C labeled UUCG-ES2 (green/purple) with ^15^N(U38) site-labeled TAR (blue) and TAR-C24U25A35 (red) in the A) absence and B) presence of 1 mM Mg^2+^. The ES2 U38-N3H3 resonance is absent in unmodified TAR and clearly visible in TAR-C24U25A35. Full [^15^N,^1^H] 2D HSQC spectra of UUCG-ES2 are shown in Figure S7.

## Discussion

Our results show that beyond biasing the local sugar pucker equilibrium, Nm can have broader effects on RNA secondary structural ensembles by preferentially stabilizing and prolonging the lifetime of conformations where Nm-modified residues adopt a paired helical conformation. Interestingly, in the U2-U6 snRNA complex, Nm modifications stabilize a four-way junction where the Nm-modified residues are paired relative to a three-way junction where they are unpaired (19), suggesting a similar mechanism to the one uncovered in our studies. Given the prevalence of secondary structural transitions during the folding, assembly, and function of both the spliceosome (95) and ribosome (96), and given the high abundance of Nm at functionally important non-canonical motifs, there may be analogous roles for Nm that involve the modulation of secondary structural equilibria in these and other Nm-modified RNAs. The preferential stabilization of paired conformations uncovered in this work should provide a useful framework for investigating such roles for Nm in RNA biology.

Nm also expands the toolbox of modifications currently available for studying RNA ESs (55). Point substitution mutations and nucleobase modifications that render an ES the dominant conformation have proven critically important for visualizing the structures of RNA ESs as well as for characterizing their functional properties (55,66,67,97-99) and testing their validity as drug targets (64). Yet because the sequence is altered, ES-mutants may not fully recapitulate the properties of the wild-type ES and finding an ES-mutant is not always feasible (58,99). Any putative model for an RNA ES could be tested by introducing Nm at residues that are unpaired in the GS but are proposed to be paired in the ES and then testing for an increase in ES abundance by NMR or chemical probing (100-102). Nm is an attractive strategy for rationally modulating RNA ensembles because it preserves the sequence identify, can be used to target all four nucleotides, and because the modification is inexpensive and commercially available for *in vitro* studies. In addition, snoRNAs can be engineered to guide novel Nm modifications *in vivo* (103), and this could provide a route for studying RNA ESs *in vivo* or provide a means for altering RNA cellular activity as a therapeutic strategy.

Nm also provides a rare opportunity to gain insights into the TS of conformational exchange through Φ-value analysis, and specifically, to gain insights into whether or not a given residue is paired (C3’-*endo*) or unpaired (enriched in C2’-*endo*) in the TS. The results for ES2 in the presence of Mg^2+^ suggest an early TS2 in which all the modified residues remain unpaired and enriched in the C2’-*endo* sugar pucker as in the GS. An Nm scan across the TAR sequence combined with RD measurements will permit broader characterization of the TS structure, and help to elucidate the base pairs that might partially melt in the TS to initiate pair reshuffling and produce ES2. These structural insights into the TS2 are of great interest in the development of anti-HIV therapeutics given that the stabilization of ES2 potently inhibits the cellular activity of TAR (97) and that stabilizing the TS2 in addition to ES2 by small molecules would provide a route for accelerating production of the inhibitory ES2 conformation.

As is the case for other post-transcriptional modifications such as m^6^A (78,88,89), the impact of Nm on the TAR dynamic ensemble cannot be fully understood without an appreciation for how Mg^2+^ impacts the ensemble as well (Figure 6). Whereas Nm perturbed the GS-ES1 exchange in a largely Mg^2+^ independent manner, both the kinetics and thermodynamics of GS-ES2 exchange were strongly dependent on Mg^2+^. Based on the crystal structure of TAR bound to Ca^2+^ (91), there are no direct contacts between bound metal ions and the 2’-OH of the Nm-modified residues (91). Rather, Nm modifications could potentially destabilize conformations within the TAR GS ensemble that preferentially bind Mg^2+^. Further studies are needed to understand how this inter-play between Nm and Mg^2+^ impacts RNA folding and function.

**Figure 6.**
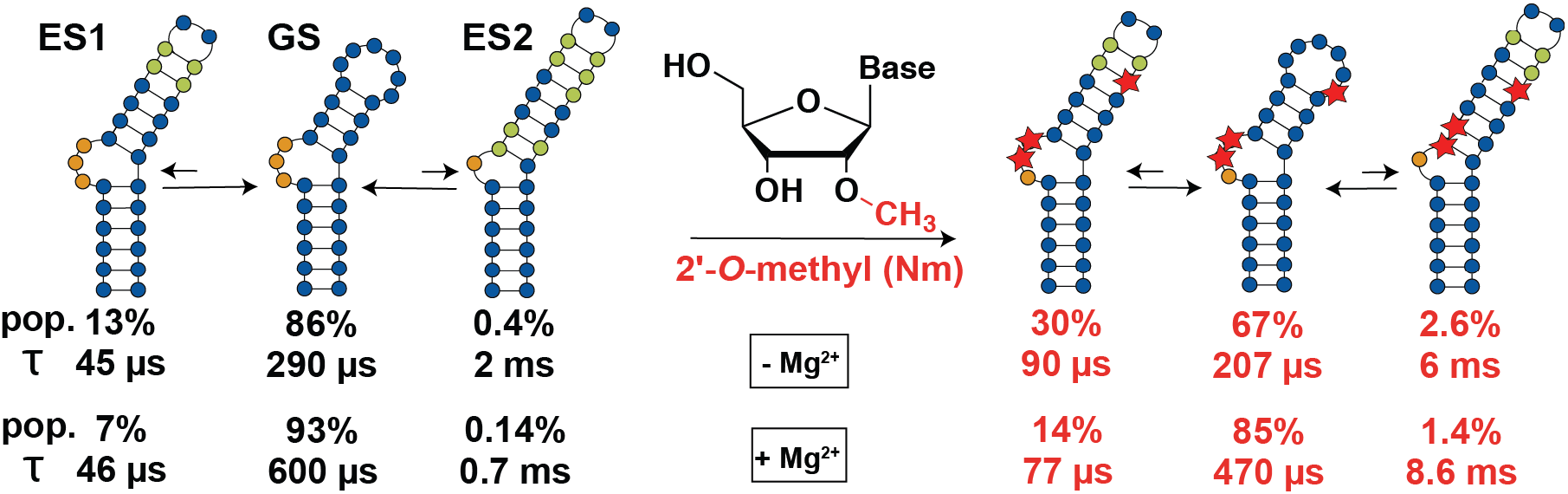
Summary of the effect of Nm on GS-ES exchange in the presence and absence of Mg^2+^. Bulge nucleotides are shown in orange. Nucleotides that transition from being unpaired in GS to paired in the ES are shown in green. Red stars indicate the Nm-modified residues. Pop and τ refer to the population and lifetime of the conformational state (GS, ES1 or ES2).

## SUPPLEMENTARY DATA

Supplementary Data are available at NAR online.

## ACKNOWLEDGEMENT

We thank the Duke Magnetic Resonance Spectroscopy Center for their technical support and resources. We also thank Atul K. Rangadurai and Megan Kelly for technical assistance and support and Dr. Richard Brennan for providing access to the UV-Vis spectrophotometer.

