## Supplementary Information for "2’-*O*-methylation alters the RNA secondary structural ensemble"

#### Supplementary Figures

**A**

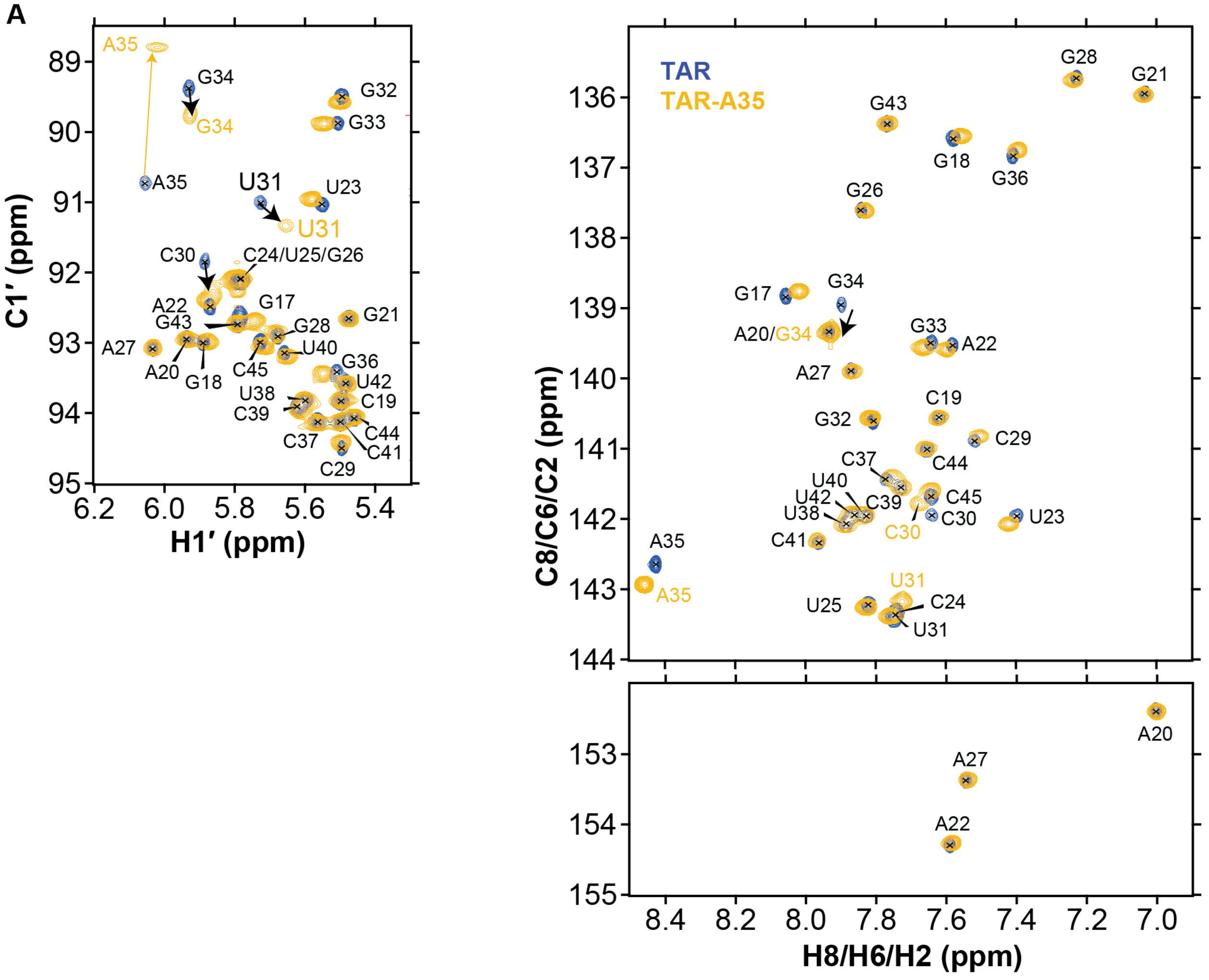

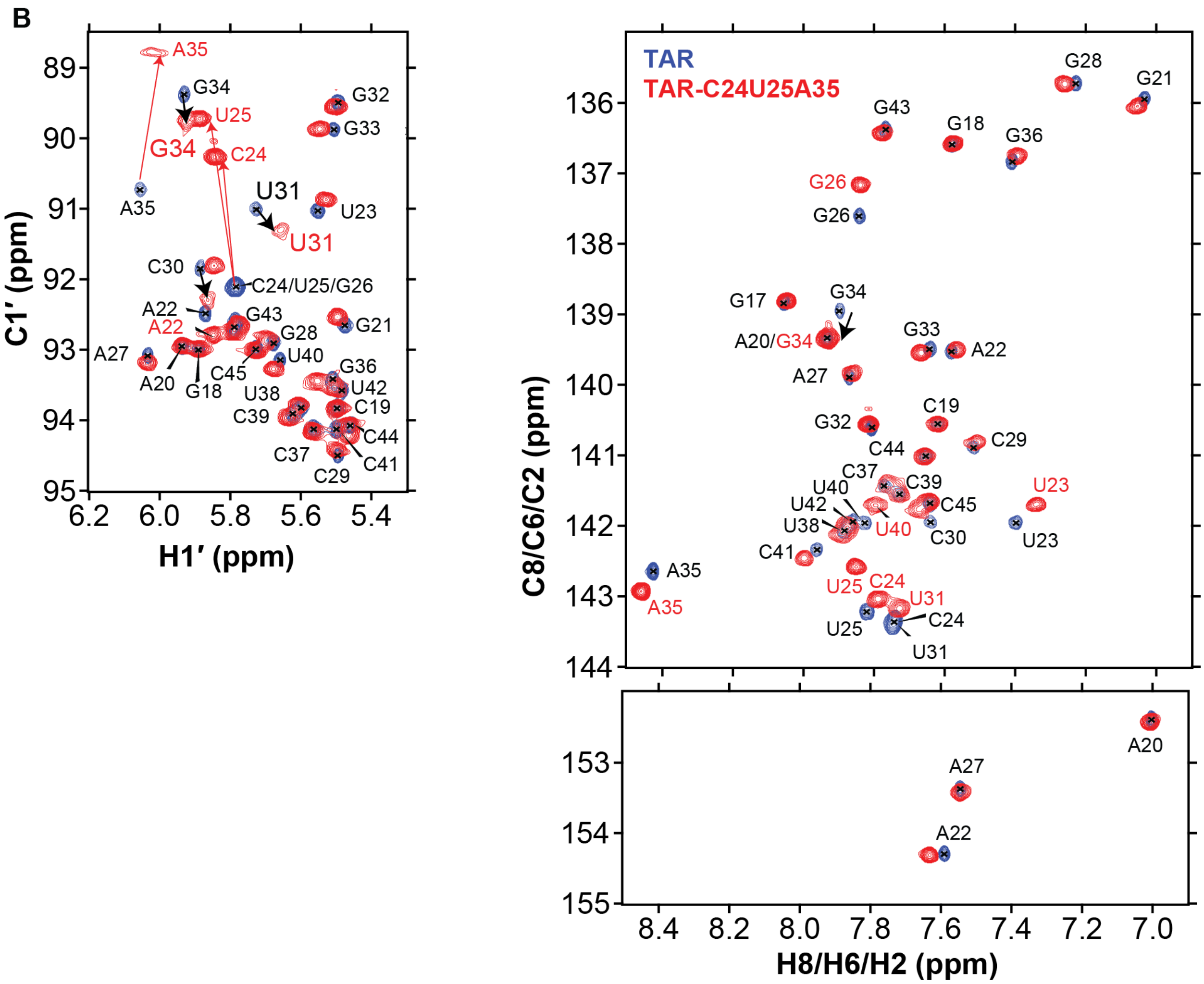


**Figure S1.** Overlay of unlabeled 2D [$^{13}$C, $^{1}$H] HSQC spectra of A) TAR-A35 (pH 6.4) and B) TAR-C24U25A35 (pH 6.4) with TAR (pH 6.4). Black arrows for the C30-C1′, U31-C1′, G34-C1′ and G34-C8 indicate CSPs towards ES1 in TAR-A35 and TAR-C24U25A35. Yellow arrow for A35-C1′ in TAR-A35 and red arrows for C24-C1′, U25-C1′ and A35-C1′ in TAR-C24U25A35 indicate the upfield shift of C1′ due to the chemical modification at the 2′ position. Sample conditions: 15 mM $NaPO_4$, 25 mM NaCl and 0.1 mM EDTA, pH 6.4, and 25 °C.

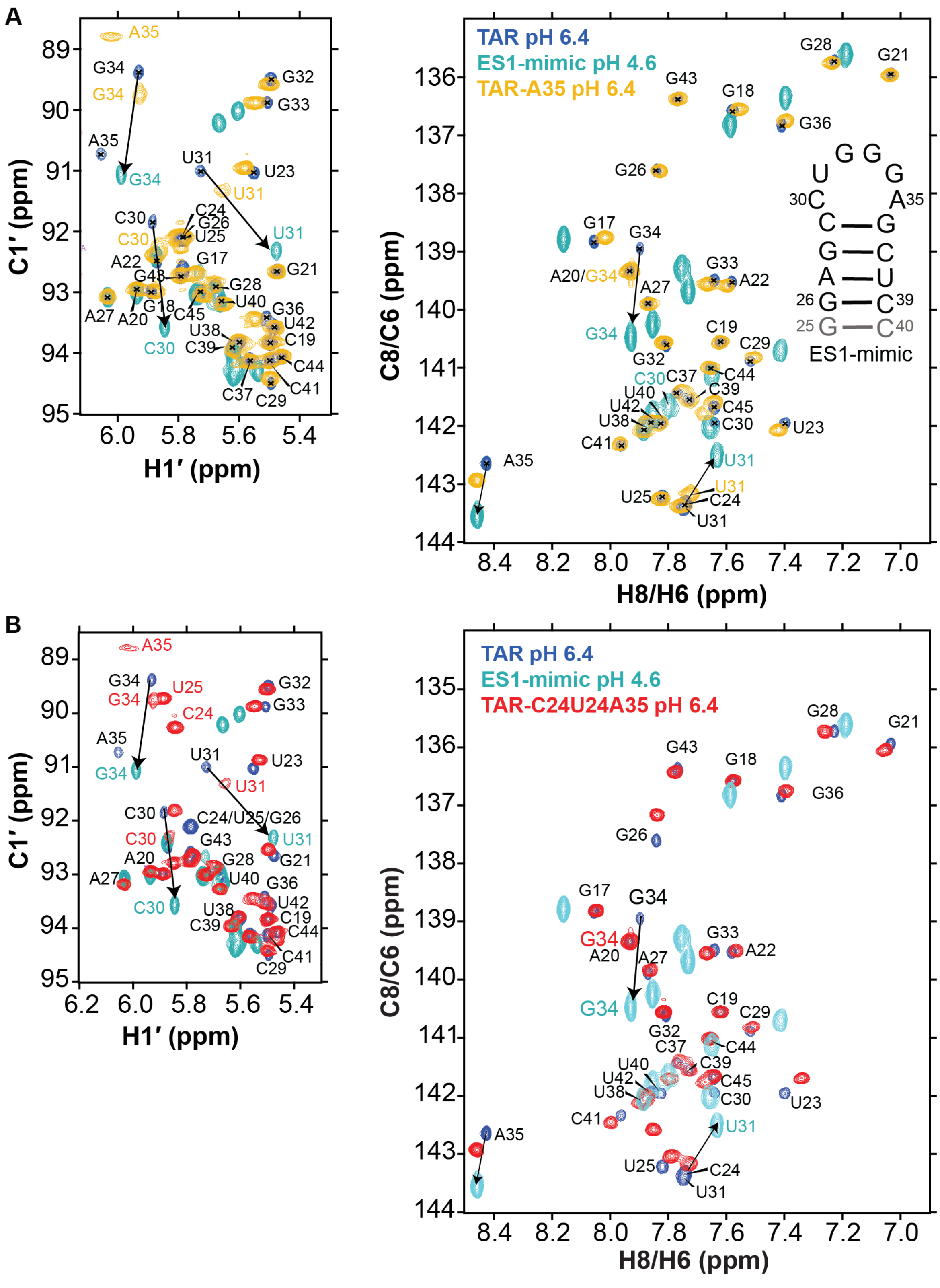


**Figure S2.** Overlay of C1′-H1′ and aromatic region of unlabeled 2D [[13]C, [1]H] HSQC spectra of TAR (pH 6.4) and a TAR ES1-mimic (pH 4.6) (1) with A) TAR-A35 (pH 6.4) and B) TAR-C24U25A35 (pH 6.4). In the previously reported TAR ES1-mimic (1), the bulge and lower helix were omitted to simplify analysis of ES1. Black arrows for the C30-C1′, U31-C1′, G34-C1′ and G34-C8 indicate a shift towards ES1 chemical shifts in TAR-A35 and TAR-C24U25A35. Sample conditions: 15 mM $NaPO_4$, 25 mM NaCl and 0.1 mM EDTA, pH 6.4, and 25 °C.

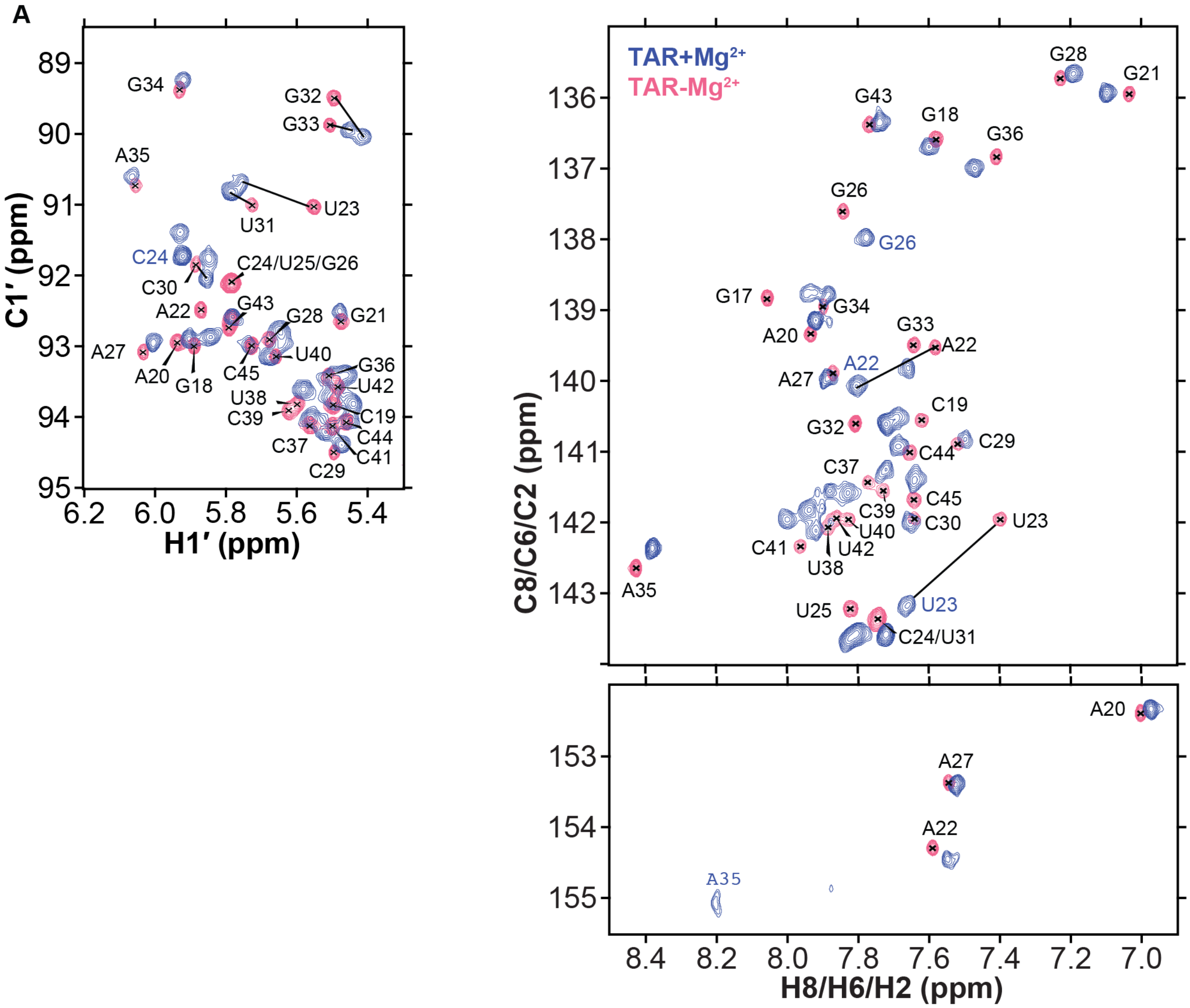

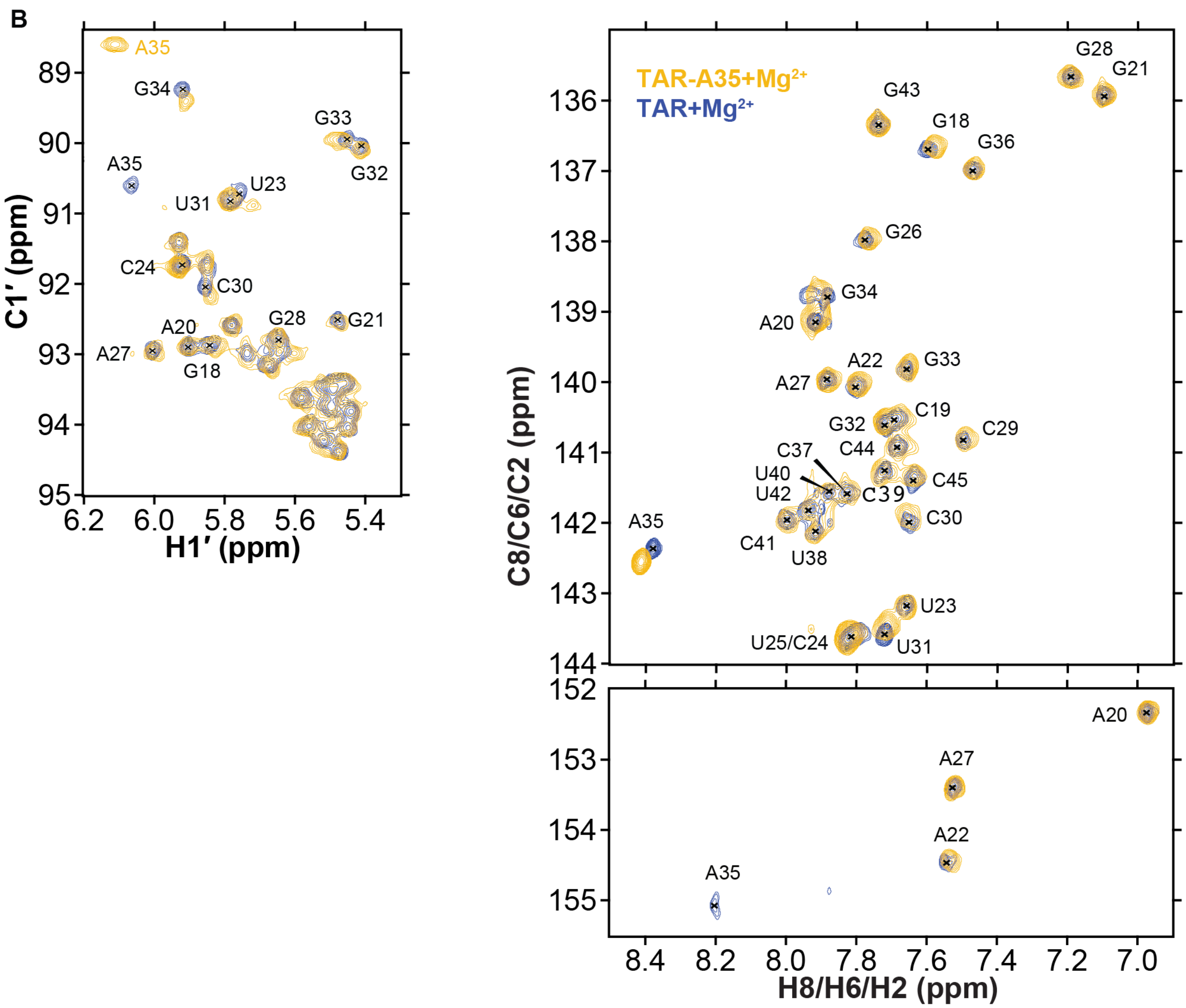

B
A35
G34
G33
G32
A35
U31
U23
C24
C30
G28
G21
A20
A27
G18
C1′ (ppm)
H1′ (ppm)
TAR-A35+Mg2+
TAR+Mg2+
G28
G21
G43
G18
G36
G26
G34
A20
A22
G33
A27
C19
G32
C29
C37
C44
U40
U42
C39
C45
C30
A35
C41
U38
U23
U25/C24
U31
A20
A27
A22
A35
C8/C6/C2 (ppm)
H8/H6/H2 (ppm)

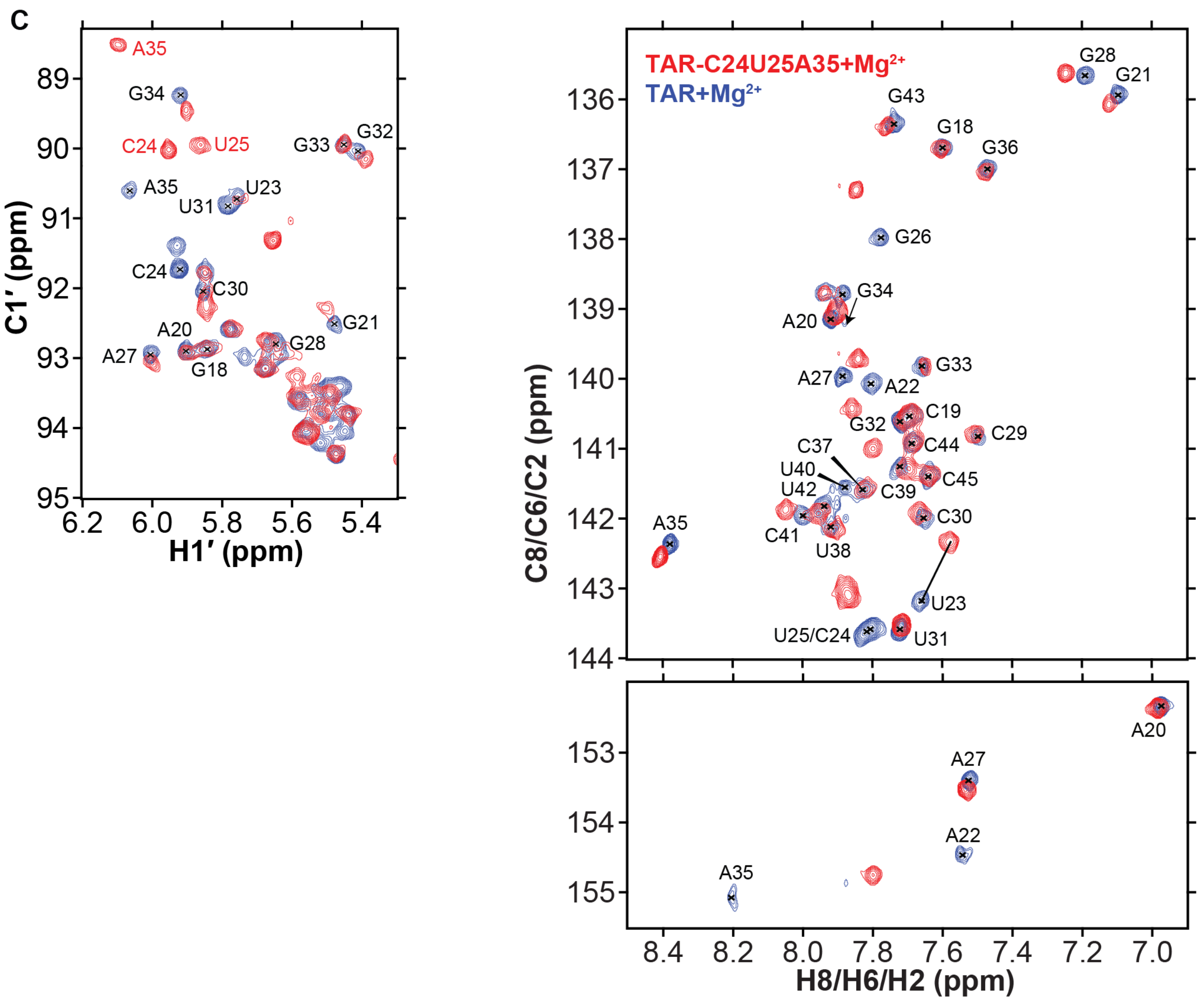
c
A35
G34
C24
U25
G33
G32
A35
U23
U31
C24
C30
G21
A20
A27
G28
G18
C1′ (ppm)
H1′ (ppm)
TAR-C24U25A35+Mg2+
TAR+Mg2+
G28
G21
G43
G18
G36
G26
G34
A20
G33
A27
A22
C19
G32
C44
C29
C37
U40
C45
U42
C39
C30
A35
C41
U38
U23
U25/C24
U31
A20
A27
A22
A35
C8/C6/C2 (ppm)
H8/H6/H2 (ppm)

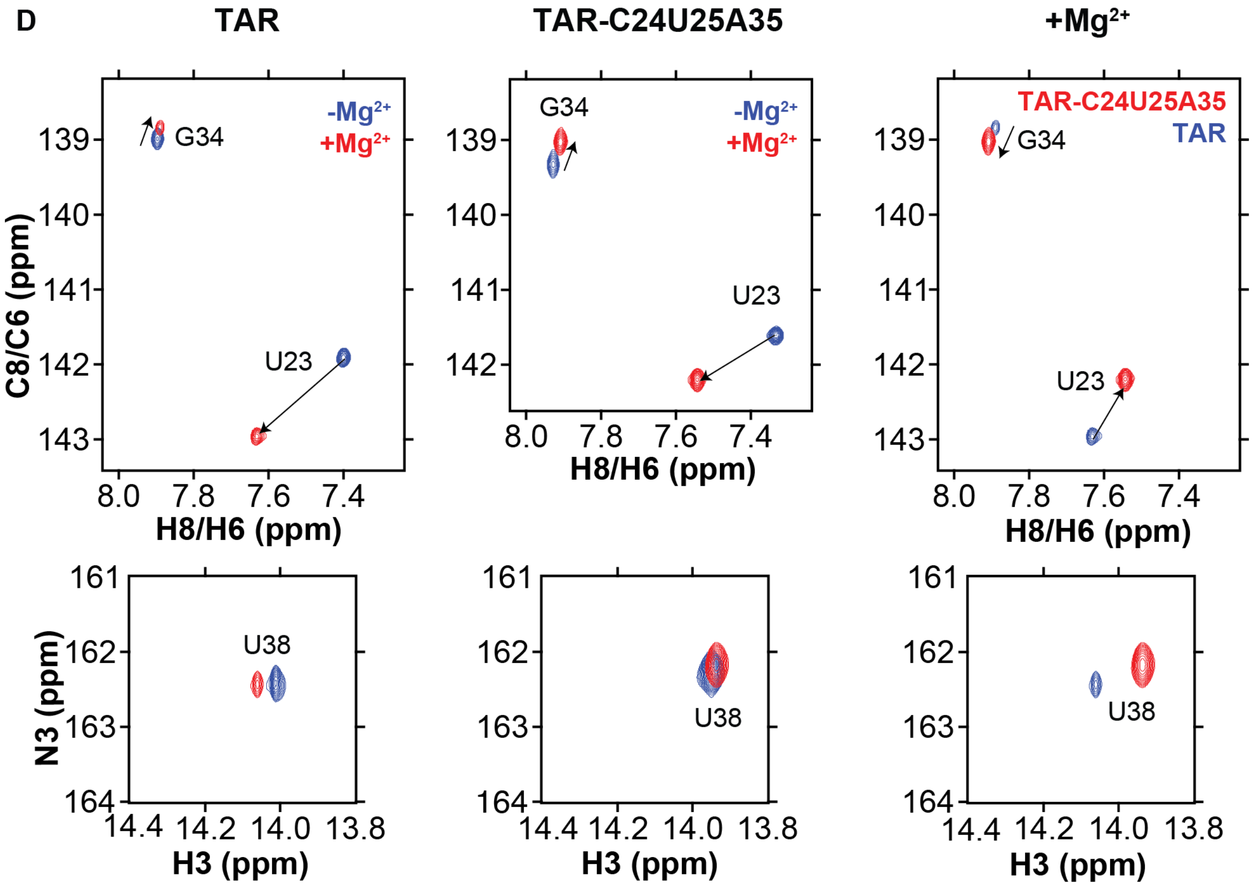


**Figure S3.** Overlay of unlabeled C1′-H1′ and aromatic region of 2D [$^{13}$C, $^{1}$H] HSQC spectra of TAR (pH 6.4) in the presence of 3 mM $Mg^{2+}$ with A) TAR (pH 6.4) in the absence of $Mg^{2+}$, B) TAR-A35 (pH 6.4) in 3 mM $Mg^{2+}$ and C) TAR-C24U25A35 (pH 6.4) in 3 mM $Mg^{2+}$. D) Overlay of site-labeled 2D [$^{13}$C, $^{1}$H] and 2D [$^{15}$N, $^{1}$H] of TAR (pH 6.4) and TAR-C24U25A35 (pH 6.4) in the presence and absence of 1 mM $Mg^{2+}$. Samples were isotopically site-labeled with $^{13}$C at G34-C8 and U23-C6 and $^{15}$N at U38-N3. Sample conditions: 15 mM $NaPO_4$, 25 mM NaCl and 0.1 mM EDTA, pH 6.4, and 25 °C.

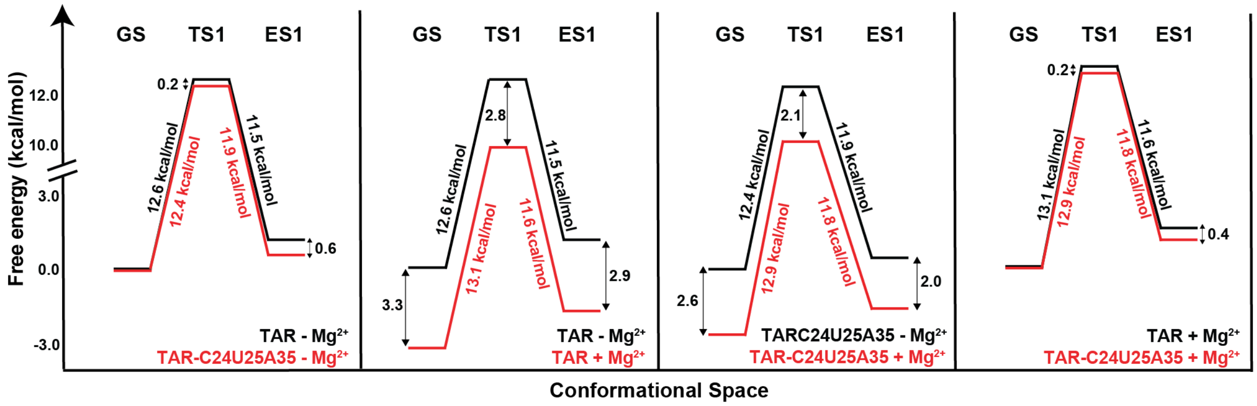


**Figure S4.** Free energy diagrams of GS-ES1 exchange.

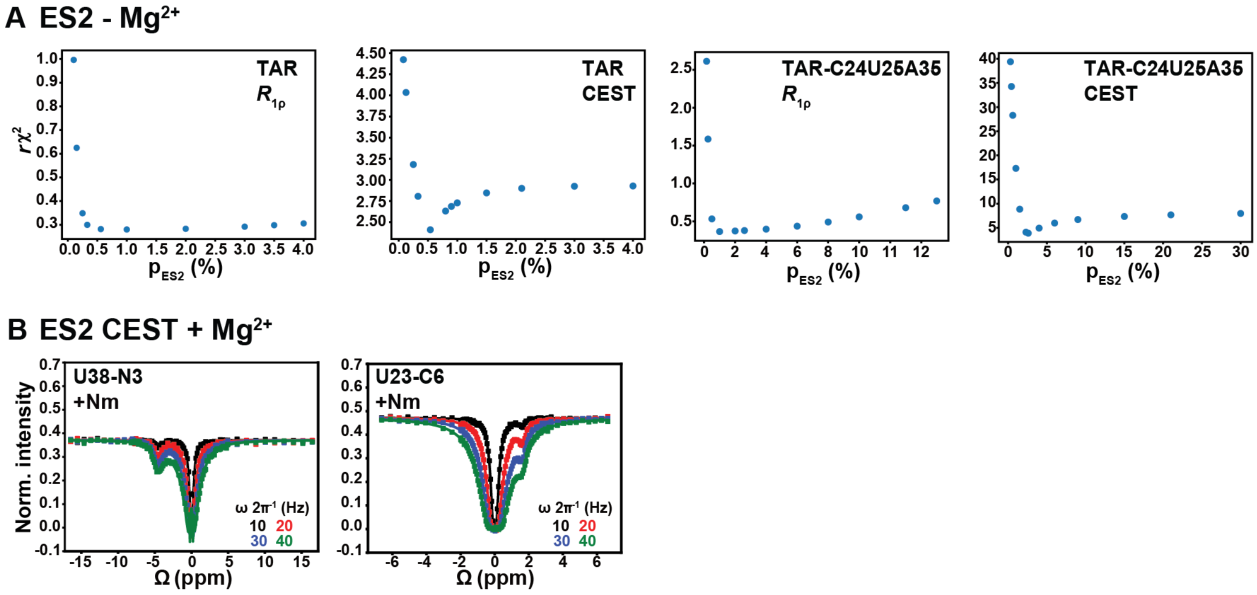


**Figure S5.** Supplementary $R_{1\rho}$ and CEST data for ES2. A) Uncertainty in exchange parameters obtained from fitting the $R_{1\rho}$ and CEST data. Shown are plots of reduced $\chi^2$ ($r\chi^2$) as a function of fixing the population of ES2 to different values for TAR and TAR-C24U25A35 in the absence of $Mg^{2+}$. B) $^{15}N$ and $^{13}C$ CEST profiles for U23-C6 and U38-N3 for TAR-C24U25A35 (pH 6.4) at 25°C in the presence of 1 mM $Mg^{2+}$. $\Omega = \omega_{RF}-\omega_{OBS}$. Spin-lock powers are color-coded.

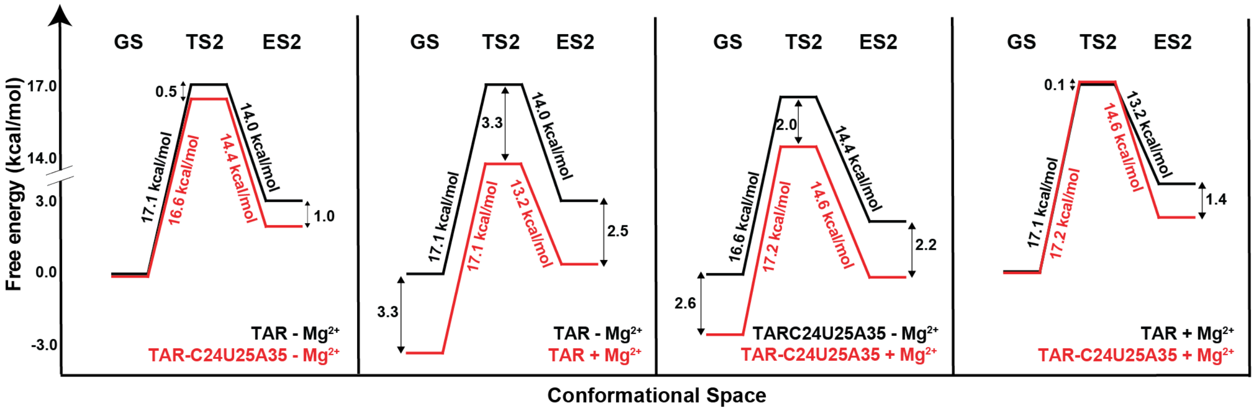


**Figure S6.** Free energy diagrams of GS-ES2 exchange.

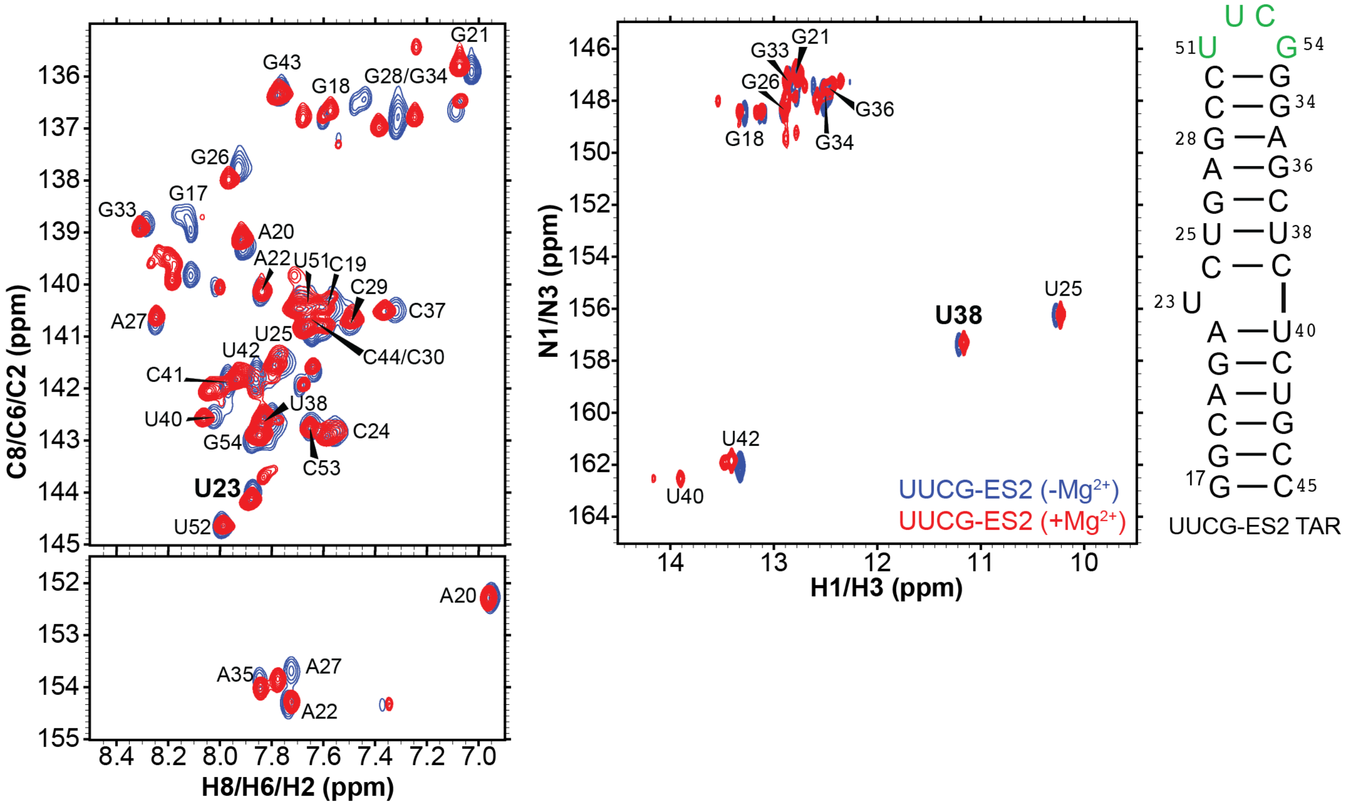


**Figure S7.** The effect of $Mg^{2+}$ on the chemical shifts of ES2 probes. Overlay of uniformly $^{13}C/^{15}N$ labeled [$^{13}C,^{1}H$] and [$^{15}N,^{1}H$] 2D HSQC spectra of UUCG-ES2 mutant (3) in the presence and absence of 1 mM $Mg^{2+}$. UUCG-ES2 stabilizes ES2 (3) as the dominant conformation through replacement of the native TAR apical loop with a UUCG loop. The addition of 1 mM $Mg^{2+}$ does not significantly affect the chemical shifts of the ES2 probes (U23-C6 and U38-N3) used in this study. Sample conditions: 25 mM NaCl, 15 mM sodium phosphate, 0.1 mM EDTA and 10% $D_2O$ at pH 6.4 and 25 °C.

### Tables

**Table S1.** Perturbations in chemical shifts of A and T/U upon 2′-*O*-methylation computed using density functional theory (DFT) calculations.

| Entry | | A-C8 | A-C2 | A-C1' | A-C2' | A-C3' | A-C4' | T/U-C6 | T/U-C1' | T/U-C2' | T/U-C3' | T/U-C4' |
|---|---|---|---|---|---|---|---|---|---|---|---|---|
| 1 | rA C3'-endo | 142.56 | 156.21 | 95.316 | 78.8973 | 70.1277 | 85.4643 | | | | | |
| 2 | rU C3'-endo | | | | | | | 147.234 | 93.9691 | 77.7646 | 69.8541 | 85.629 |
| 3 | rA C2'-endo | 140.01 | 157.01 | 94.465 | 75.158 | 77.4695 | 89.7878 | | | | | |
| 4 | rU C2'-endo | | | | | | | 145.029 | 94.6564 | 74.5493 | 78.1384 | 88.483 |
| 5 | Am, C3'-endo | 142.71 | 155.96 | 92.459 | 90.5046 | 70.39 | 85.9056 | | | | | |
| 6 | Um, C3'-endo | | | | | | | 146.916 | 91.3198 | 89.5421 | 70.1019 | 86.499 |
| 7 | Am, C2'-endo | 140.75 | 156.66 | 92.608 | 83.1558 | 77.2877 | 89.109 | | | | | |
| 8 | Um, C2'-endo | | | | | | | 145.555 | 93.7743 | 82.5949 | 77.1266 | 88.861 |

**Table S2.** Population of ES1 derived from chemical shift perturbation analysis.

| | Shift toward ES1 (%) | | | |
|---|---|---|---|---|
| | $-Mg^{2+}$ | | $+Mg^{2+}$ | |
| Resonance | TAR-A35 | TAR-C24U24A35 | TAR | TAR-C24U24A35 |
| C30C1′-H1′ | 30.5 | 34.1 | | |
| U31C1′-H1′ | 28.7 | 27.1 | | |
| U31C6-H6 | 27.5 | 27.7 | | |
| G34C1′-H1′ | 29.7 | 29.2 | | |
| G34C8-H8 | 27.0 | 26.1 | 7.3 | 14.5 |
| Average | 28.7 | 28.8 | | |
| STDEV | 1.5 | 3.1 | | |

**Table S3.** Exchange parameters for ES1 from fitting $R_{1\rho}$ data for G34-C8 at 25 °C.

| | $R_1$ (Hz) | $R_2$ (Hz) | $p_{ES1}$ (%) | Δω (ppm) | $k_{ex}$ ($s^{-1}$) | $k_1$ ($s^{-1}$) | $k_{-1}$ ($s^{-1}$) | τ (μs) | ΔG° (kcal/mol) |
|---|---|---|---|---|---|---|---|---|---|
| 1) TAR $-Mg^{2+}$ from (1) | 2.3 ± 0.1 | 22.3 ± 1.2 | 13.0 ± 2.0 | 2.6 ± 0.2 | 25807 ± 716 | 3436 ± 509 | 22371 ± 1771 | 45 ± 4 | 1.1 ± 0.1 |
| 2) TAR $-Mg^{2+}$ | 3.4 ± 0.3 | 28.9 ± 5.6 | 21.9 ± 10.0 | 2.6 ± 0.5 | 32248 ± 4214 | 7055 ± 2839 | 25193 ± 4248 | 40 ± 7 | 0.75 ± 0.3 |
| 3) TAR $-Mg^{2+}$ | 3.3 ± 0.2 | 31.3 ± 4.4 | 13* | 3.0 ± 0.4 | 30531 ± 3484 | 3969 ± 3521 | 26562 ± 4624 | 38 ± 7 | 1.1 |
| 4) TAR-C24U25A35 $-Mg^{2+}$ | 1.2 ± 0.4 | 32.6 ± 2.1 | 30.3 ± 6.0 | 2.4 ± 0.2 | 15874 ± 613 | 4816 ± 752 | 11059 ± 848 | 90 ± 7 | 0.5 ± 0.1 |
| 5) TAR-C24U25A35 $-Mg^{2+}$ | 1.2 ± 0.4 | 32.7 ± 2.1 | 29** | 2.4 ± 0.1 | 15852 ± 613 | 4597 ± 3530 | 11255 ± 3553 | 89 ± 28 | 0.5** |
| 6) TAR +1 mM $Mg^{2+}$ | 2.6 ± 0.2 | 37.3 ± 2.6 | 7.0** | 2.9 ± 0.3 | 23516 ± 2685 | 1646 ± 1547 | 21870 ± 2932 | 46 ± 6 | 1.5** |
| 7) TAR-C24U25A35 +1 mM $Mg^{2+}$ | 2.1 ± 0.2 | 37.2 ± 1.1 | 14.1 ± 2.8 | 2.5 ± 0.2 | 15098 ± 487 | 2122 ± 372 | 12976 ± 556 | 77 ± 3 | 1.1 ± 0.1 |

*values fixed while fitting the data, these values are obtained from previously published results (1)

**$p_{ES1}$ from chemical shift perturbation analysis in 2D HSQC NMR and the corresponding ΔG° is calculated based on the following equation: $RTln[p_{ES1}/(1-p_{ES1})]$

**Table S4.** Free energy values calculated for TS1 and TS2.

| ES | Condition | Δ‡G° kcal/mol |
|---|---|---|
| ES1 | TAR | 12.6 ± 0.1 |
| | TAR-C24U25A35 | 12.4 ± 0.1 |
| | TAR + $Mg^{2+}$ | 13.1 ± 0.6 |
| | TAR-C24U25A35 + $Mg^{2+}$ | 12.9 ± 0.1 |
| ES2 | TAR | 17.06 ± 0.04 |
| | TAR-C24U25A35 | 16.56 ± 0.01 |
| | TAR + $Mg^{2+}$ | 17.06 ± 0.14 |
| | TAR-C24U25A35 + $Mg^{2+}$ | 17.15 ± 0.03 |

**Table S5.** Phi-values for Nm-modified TAR in the absence of presence of $Mg^{2+}$.

| ES | Sample | Φ |
|---|---|---|
| ES1 | TAR-C24U25A35 | 0.33 ± 0.25 |
| | TAR-C24U25A35 + $Mg^{2+}$ | 0.5 ± 1.6* |
| ES2 | TAR-C24U25A35 | 0.53 ± 0.04 |
| | TAR-C24U25A35 + $Mg^{2+}$ | -0.07 ± 0.10* |

*Phi-value calculated for TAR-C24U25A35 in the presence of $Mg^{2+}$ with TAR in $Mg^{2+}$ as a reference.

**Table S6A.** Comparison of ES2 exchange parameters for TAR and TAR-C24U25A35 obtained from fitting CEST and $R_{1\rho}$ data in the absence of $Mg^{2+}$.

| | R1 (Hz) | R2 (Hz) | $p_{ES2}$ (%) | Δω (ppm) | $k_{ex}$ ($s^{-1}$) | $k_1$ ($s^{-1}$) | $k_{-1}$ ($s^{-1}$) | τ (ms) | ΔG° (kcal/mol) |
|---|---|---|---|---|---|---|---|---|---|
| TAR U23-C6 from (3) | 2.50 ± 0.04 | 30.7 ± 0.1 | 0.40 ± 0.05 | 2.3 ± 0.1 | 474 ± 69 | 1.9 ± 0.4 | 472 ± 70 | 2.1 ± 0.3 | 3.26 ± 0.07 |
| TAR U38-N3 from (3) | 1.40 ± 0.03 | 6.20 ± 0.03 | | -5.2 ± 0.2 | | | | | |
| TAR CEST (U23-C6) | 2.702 ± 0.004 | 30.38 ± 0.19 | 0.54 ± 0.01 | 2.07 ± 0.02 | 356 ± 23 | 1.92 ± 0.13 | 354 ± 23 | 2.8 ± 0.2 | 3.09 ± 0.01 |
| TAR CEST (U38-N3) | 3.111 ± 0.005 | 9.90 ± 0.13 | | -5.12 ± 0.06 | | | | | |
| TAR RD (U23-C6) | 2.71 ± 0.07 | 30.07 ± 0.09 | 0.54* | 2.32 ± 0.13 | 431 ± 15 | 2.3 ± 2.3 | 429 ± 15 | 2.33 ± 0.08 | |
| TAR RD (U38-N3) | 2.03 ± 0.02 | 6.29 ± 0.02 | | -4.89 ± 0.13 | | | | | |
| TAR-C24U25A35 CEST (U23-C6) | 2.538 ± 0.003 | 31.15 ± 0.14 | 2.60 ± 0.03 | 2.142 ± 0.003 | 171 ± 3 | 4.45 ± 0.09 | 167 ± 3 | 6.0 ± 0.1 | 2.15 ± 0.01 |
| TAR-C24U25A35 CEST (U38-N3) | 3.458 ± 0.004 | 10.21 ± 0.08 | | -4.877 ± 0.009 | | | | | |
| TAR-C24U25A35 RD (U23-C6) | 2.9 ± 0.1 | 33.53 ± 0.13 | 2.6* | 2.25 ± 0.08 | 174 ± 5 | 4.5 ± 4.4 | 170 ± 6 | 5.9 ± 0.2 | |
| TAR-C24U25A35 RD (U38-N3) | 2.15 ± 0.03 | 6.68 ± 0.03 | | -4.56 ± 0.09 | | | | | |

* $R_{1\rho}$ RD data was fit by fixing the population to the values determined by CEST.

**Table S6B.** Comparison of ES2 exchange parameters for TAR and TAR-C24U25A35 in the presence of 1 mM $Mg^{2+}$ obtained from fitting CEST and $R_{1\rho}$ data.

| | R1 (Hz) | R2 (Hz) | $p_{ES2}$ (%) | Δω (ppm) | $k_{ex}$ ($s^{-1}$) | $k_1$ ($s^{-1}$) | $k_{-1}$ ($s^{-1}$) | τ (ms) | ΔG° (kcal/mol) |
|---|---|---|---|---|---|---|---|---|---|
| TAR RD (U38-N3) | 2.02 ± 0.03 | 6.62 ± 0.05 | 0.14 ± 0.02 | -4.4 ± 0.5 | 1367 ± 267 | 1.92 ± 0.47 | 1365 ± 266 | 0.73 ± 0.14 | 3.89 ± 0.19 |
| TAR-C24U25A35 CEST (U23-C6) | 2.459 ± 0.003 | 39.25 ± 0.17 | 1.39 ± 0.03 | 1.631 ± 0.004 | 118 ± 4 | 1.65 ± 0.07 | 116 ± 4 | 8.6 ± 0.3 | 2.52 ± 0.01 |
| TAR-C24U25A35 CEST (U38-N3) | 3.255 ± 0.003 | 10.49 ± 0.07 | | -4.66 ± 0.01 | | | | | |
| TAR-C24U25A35 RD (U23-C6) | 2.6 ± 0.1 | 41.66 ± 0.13 | 1.4* | 2.0 ± 0.2 | 117 ± 5 | 1.6 ± 1.6 | 116 ± 5 | 8.6 ± 0.4 | |
| TAR-C24U25A35 RD (U38-N3) | 1.96 ± 0.02 | 7.14 ± 0.02 | | -4.43 ± 0.15 | | | | | |

* $R_{1\rho}$ RD data for TAR-C24U25A35 was fit by fixing the population to the values determined by CEST.

**Table S7.** Spin-lock power and offsets used in the $R_{1\rho}$ measurements.

| construct | Nuclei | **[spin-lock power] {offset frequencies}/[$\omega_1$ $2\pi^{-1}(s^{-1})$] {$\Omega$ $2\pi^{-1}(s^{-1})$}** |
|---|---|---|
| TAR (-$Mg^{2+}$) | U23-C6 | [50, 100, 150, 200, 250, 300, 350, 400, 500, 600, 700, 800, 900, 1000, 1200, 1400, 1600, 1800, 2000, 2500] {0} |
| | | [200] {-800, -700, -600, -540, -480, -420, -390, -360, -330, -300, -270, -240, -210, -180, -150, -120, -90, -60, -30, 30, 60, 90, 120, 150, 180, 210, 240, 270, 300, 330, 360, 390, 420, 480, 540, 600, 700, 800} |
| | | [400] {-1500, -1200, -1000, -900, -800, -700, -600, -550, -500, -450, -400, -350, -300, -250, -200, -150, -100, -50, 50, 100, 150, 200, 250, 300, 350, 400, 450, 500, 550, 600, 700, 800, 900, 1000, 1200, 1500} |
| | | [600] {-1600, -1400, -1200, -1000, -800, -650, -550, -450, -375, -300, -225, -150, -75, 75, 150, 225, 300, 375, 450, 550, 650, 800, 1000, 1200, 1400, 1600} |
| | | [1000] {-1700, -1500, -1300, -1100, -900, -700, -500, -400, -300, -200, -100, 100, 200, 300, 400, 500, 700, 900, 1100, 1300, 1500, 1700} |
| | U38-N3 | [100, 150, 200, 250, 300, 350, 400, 450, 500, 500, 600, 700, 800, 900, 1000, 1200, 1400, 1600, 1800, 2000] {0} |
| | | [200] {-500, -400, -350, -300, -250, -225, -200, -180, -160, -140, -120, -100, -80, -60, -40, -20, 20, 40, 60, 80, 100, 120, 140, 160, 180, 200, 225, 250, 300, 350, 400, 500} |
| | | [400] {-1000, -800, -700, -600, -500, -400, -350, -300, -250, -210, -180, -150, -120, -90, -60, -30, 30, 60, 90, 120, 150, 180, 210, 250, 300, 350, 400, 500, 600, 700, 800, 1000} |
| | | [600] {-1600, -1400, -1200, -1000, -800, -600, -500, -400, -300, -240, -200, -160, -120, -80, -40, 40, 80, 120, 160, 200, 240, 300, 400, 500, 600, 800, 1000, 1200, 1400, 1600} |
| | | [1000] {-3000, -2500, -2000, -1500, -1300, -1100, -900, -700, -600, -500, -400, -300, -200, -100, 100, 200, 300, 400, 500, 600, 700, 900, 1100, 1300, 1500, 2000, 2500, 3000} |
| TAR-C24U25A35 (-$Mg^{2+}$) | U23-C6 | [200, 250, 300, 350, 400, 500, 600, 700, 800, 900, 1000, 1200, 1400, 1600, 1800, 2000, 2500] {0} |
| | | [200] {-800, -700, -600, -540, -480, -420, -390, -360, -330, -300, -270, -240, -210, -180, -150, -120, -90, -60, -30, 30, 60, 90, 120, 150, 180, 210, 240, 270, 300, 330, 360, 390, 420, 480, 540, 600, 700, 800} |
| | | [400] {-1500, -1200, -1000, -900, -800, -700, -600, -550, -500, -450, -400, -350, -300, -250, -200, -150, -100, -50, 50, 100, 150, 200, 250, 300, 350, 400, 450, 500, 550, 600, 700, 800, 900, 1000, 1200, 1500} |
| | | [600] {-1600, -1400, -1200, -1000, -800, -650, -550, -450, -375, -300, -225, -150, -75, 75, 150, 225, 300, 375, 450, 550, 650, 800, 1000, 1200, 1400, 1600} |
| | | [1000] {-1700, -1500, -1300, -1100, -900, -700, -500, -400, -300, -200, -100, 100, 200, 300, 400, 500, 700, 900, 1100, 1300, 1500, 1700} |
| | U38-N3 | [100, 150, 200, 250, 300, 350, 400, 450, 500, 500, 600, 700, 800, 900, 1000, 1200, 1400, 1600, 1800, 2000] {0} |
| | | [200] {-500, -400, -350, -300, -250, -225, -200, -180, -160, -140, -120, -100, -80, -60, -40, -20, 20, 40, 60, 80, 100, 120, 140, 160, 180, 200, 225, 250, 300, 350, 400, 500} |
| | | [400] {-1000, -800, -700, -600, -500, -400, -350, -300, -250, -210, -180, -150, -120, -90, -60, -30, 30, 60, 90, 120, 150, 180, 210, 250, 300, 350, 400, 500, 600, 700, 800, 1000} |
| | | [600] {-1600, -1400, -1200, -1000, -800, -600, -500, -400, -300, -240, -200, -160, -120, -80, -40, 40, 80, 120, 160, 200, 240, 300, 400, 500, 600, 800, 1000, 1200, 1400, 1600} |
| | | [1000] {-3000, -2500, -2000, -1500, -1300, -1100, -900, -700, -600, -500, -400, -300, -200, -100, 100, 200, 300, 400, 500, 600, 700, 900, 1100, 1300, 1500, 2000, 2500, 3000} |
| TAR (+$Mg^{2+}$) | U38-N3 | [100, 150, 200, 250, 300, 350, 400, 450, 500, 500, 600, 700, 800, 900, 1000, 1200, 1400, 1600, 1800, 2000] {0} |

| | | |
|---|---|---|
| | | [200] {-500, -400, -350, -300, -250, -225, -200, -180, -160, -140, -120, -100, -80, -60, -40, -20, 20, 40, 60, 80, 100, 120, 140, 160, 180, 200, 225, 250, 300, 350, 400, 500} |
| | | [400] {-1000, -800, -700, -600, -500, -400, -350, -300, -250, -210, -180, -150, -120, -90, -60, -30, 30, 60, 90, 120, 150, 180, 210, 250, 300, 350, 400, 500, 600, 700, 800, 1000} |
| | | [600] {-1600, -1400, -1200, -1000, -800, -600, -500, -400, -300, -240, -200, -160, -120, -80, -40, 40, 80, 120, 160, 200, 240, 300, 400, 500, 600, 800, 1000, 1200, 1400, 1600} |
| | | [1000] {-3000, -2500, -2000, -1500, -1300, -1100, -900, -700, -600, -500, -400, -300, -200, -100, 100, 200, 300, 400, 500, 600, 700, 900, 1100, 1300, 1500, 2000, 2500, 3000} |
| TAR-C24U25A35 (+$Mg^{2+}$) | U23-C6 | [50, 100, 150, 200, 250, 300, 350, 400, 500, 600, 700, 800, 900, 1000, 1200, 1400, 1600, 1800, 2000, 2500] {0} |
| | | [200] {-800, -700, -600, -540, -480, -420, -390, -360, -330, -300, -270, -240, -210, -180, -150, -120, -90, -60, -30, 30, 60, 90, 120, 150, 180, 210, 240, 270, 300, 330, 360, 390, 420, 480, 540, 600, 700, 800} |
| | | [400] {-1500, -1200, -1000, -900, -800, -700, -600, -550, -500, -450, -400, -350, -300, -250, -200, -150, -100, -50, 50, 100, 150, 200, 250, 300, 350, 400, 450, 500, 550, 600, 700, 800, 900, 1000, 1200, 1500} |
| | | [600] {-1600, -1400, -1200, -1000, -800, -650, -550, -450, -375, -300, -225, -150, -75, 75, 150, 225, 300, 375, 450, 550, 650, 800, 1000, 1200, 1400, 1600} |
| | | [1000] {-1700, -1500, -1300, -1100, -900, -700, -500, -400, -300, -200, -100, 100, 200, 300, 400, 500, 700, 900, 1100, 1300, 1500, 1700} |
| | U38-N3 | [100, 150, 200, 250, 300, 350, 400, 450, 500, 500, 600, 700, 800, 900, 1000, 1200, 1400, 1600, 1800, 2000] {0} |
| | | [200] {-500, -400, -350, -300, -250, -225, -200, -180, -160, -140, -120, -100, -80, -60, -40, -20, 20, 40, 60, 80, 100, 120, 140, 160, 180, 200, 225, 250, 300, 350, 400, 500} |
| | | [400] {-1000, -800, -700, -600, -500, -400, -350, -300, -250, -210, -180, -150, -120, -90, -60, -30, 30, 60, 90, 120, 150, 180, 210, 250, 300, 350, 400, 500, 600, 700, 800, 1000} |
| | | [600] {-1600, -1400, -1200, -1000, -800, -600, -500, -400, -300, -240, -200, -160, -120, -80, -40, 40, 80, 120, 160, 200, 240, 300, 400, 500, 600, 800, 1000, 1200, 1400, 1600} |
| | | [1000] {-3000, -2500, -2000, -1500, -1300, -1100, -900, -700, -600, -500, -400, -300, -200, -100, 100, 200, 300, 400, 500, 600, 700, 900, 1100, 1300, 1500, 2000, 2500, 3000} |

**Table S8.** Spin-lock power and offsets used in the CEST measurements.

| construct | Nuclei | **[spin-lock power] {offset frequencies}** |
|---|---|---|
| | | **[$\omega_1$ $2\pi^{-1}(s^{-1})$] {$\Omega$ $2\pi^{-1}(s^{-1})$}** |
| TAR ($-Mg^{2+}$) | U23-C6 | [10] {-1000.0, -922.2, -844.4, -766.7, -688.9, -611.1, -533.3, -455.6, -377.8, -300.0, -300.0, -289.9, -279.7, -269.6, -259.5, -249.4, -239.2, -229.1, -219.0, -208.9, -198.7, -188.6, -178.5, -168.4, -158.2, -148.1, -138.0, -127.8, -117.7, -107.6, -97.5, -87.3, -77.2, -67.1, -57.0, -46.8, -36.7, -26.6, -16.5, -6.3, 3.8, 13.9, 24.1, 34.2, 44.3, 54.4, 64.6, 74.7, 84.8, 94.9, 105.1, 115.2, 125.3, 135.4, 145.6, 155.7, 165.8, 175.9, 186.1, 196.2, 206.3, 216.5, 226.6, 236.7, 246.8, 257.0, 267.1, 277.2, 287.3, 297.5, 307.6, 317.7, 327.8, 338.0, 348.1, 358.2, 368.4, 378.5, 388.6, 398.7, 408.9, 419.0, 429.1, 439.2, 449.4, 459.5, 469.6, 479.7, 489.9, 500.0, 500.0, 555.6, 611.1, 666.7, 722.2, 777.8, 833.3, 888.9, 944.4, 1000.0} |
| | | [20] {-1000.0, -922.2, -844.4, -766.7, -688.9, -611.1, -533.3, -455.6, -377.8, -300.0, -300.0, -289.9, -279.7, -269.6, -259.5, -249.4, -239.2, -229.1, -219.0, -208.9, -198.7, -188.6, -178.5, -168.4, -158.2, -148.1, -138.0, -127.8, -117.7, -107.6, -97.5, -87.3, -77.2, -67.1, -57.0, -46.8, -36.7, -26.6, -16.5, -6.3, 3.8, 13.9, 24.1, 34.2, 44.3, 54.4, 64.6, 74.7, 84.8, 94.9, 105.1, 115.2, 125.3, 135.4, 145.6, 155.7, 165.8, 175.9, 186.1, 196.2, 206.3, 216.5, 226.6, 236.7, 246.8, 257.0, 267.1, 277.2, 287.3, 297.5, 307.6, 317.7, 327.8, 338.0, 348.1, 358.2, 368.4, 378.5, 388.6, 398.7, 408.9, 419.0, 429.1, 439.2, 449.4, 459.5, 469.6, 479.7, 489.9, 500.0, 500.0, 555.6, 611.1, 666.7, 722.2, 777.8, 833.3, 888.9, 944.4, 1000.0} |
| | | [30] {-1000.0, -922.2, -844.4, -766.7, -688.9, -611.1, -533.3, -455.6, -377.8, -300.0, -300.0, -289.9, -279.7, -269.6, -259.5, -249.4, -239.2, -229.1, -219.0, -208.9, -198.7, -188.6, -178.5, -168.4, -158.2, -148.1, -138.0, -127.8, -117.7, -107.6, -97.5, -87.3, -77.2, -67.1, -57.0, -46.8, -36.7, -26.6, -16.5, -6.3, 3.8, 13.9, 24.1, 34.2, 44.3, 54.4, 64.6, 74.7, 84.8, 94.9, 105.1, 115.2, 125.3, 135.4, 145.6, 155.7, 165.8, 175.9, 186.1, 196.2, 206.3, 216.5, 226.6, 236.7, 246.8, 257.0, 267.1, 277.2, 287.3, 297.5, 307.6, 317.7, 327.8, 338.0, 348.1, 358.2, 368.4, 378.5, 388.6, 398.7, 408.9, 419.0, 429.1, 439.2, 449.4, 459.5, 469.6, 479.7, 489.9, 500.0, 500.0, 555.6, 611.1, 666.7, 722.2, 777.8, 833.3, 888.9, 944.4, 1000.0} |
| | | [40] {-1000.0, -922.2, -844.4, -766.7, -688.9, -611.1, -533.3, -455.6, -377.8, -300.0, -300.0, -289.9, -279.7, -269.6, -259.5, -249.4, -239.2, -229.1, -219.0, -208.9, -198.7, -188.6, -178.5, -168.4, -158.2, -148.1, -138.0, -127.8, -117.7, -107.6, -97.5, -87.3, -77.2, -67.1, -57.0, -46.8, -36.7, -26.6, -16.5, -6.3, 3.8, 13.9, 24.1, 34.2, 44.3, 54.4, 64.6, 74.7, 84.8, 94.9, 105.1, 115.2, 125.3, 135.4, 145.6, 155.7, 165.8, 175.9, 186.1, 196.2, 206.3, 216.5, 226.6, 236.7, 246.8, 257.0, 267.1, 277.2, 287.3, 297.5, 307.6, 317.7, 327.8, 338.0, 348.1, 358.2, 368.4, 378.5, 388.6, 398.7, 408.9, 419.0, 429.1, 439.2, 449.4, 459.5, 469.6, 479.7, 489.9, 500.0, 500.0, 555.6, 611.1, 666.7, 722.2, 777.8, 833.3, 888.9, 944.4, 1000.0} |
| TAR ($-Mg^{2+}$) | U38-N3 | [10] {-800.0, -766.7, -733.3, -700.0, -666.7, -633.3, -600.0, -566.7, -533.3, -500.0, -500.0, -489.9, -479.7, -469.6, -459.5, -449.4, -439.2, -429.1, -419.0, -408.9, -398.7, -388.6, -378.5, -368.4, -358.2, -348.1, -338.0, -327.8, -317.7, -307.6, -297.5, -287.3, -277.2, -267.1, -257.0, -246.8, -236.7, -226.6, -216.5, -206.3, -196.2, -186.1, -175.9, -165.8, -155.7, -145.6, -135.4, -125.3, -115.2, -105.1, -94.9, -84.8, -74.7, -64.6, -54.4, -44.3, -34.2, -24.1, -13.9, -3.8, 6.3, 16.5, 26.6, 36.7, 46.8, 57.0, 67.1, 77.2, 87.3, 97.5, 107.6, 117.7, 127.8, 138.0, 148.1, 158.2, 168.4, 178.5, 188.6, 198.7, 208.9, 219.0, 229.1, 239.2, 249.4, 259.5, 269.6, 279.7, 289.9, 300.0, 300.0, 333.3, 366.7, 400.0, 433.3, 466.7, 500.0, 533.3, 566.7, 600.0} |
| | | [20] {-800.0, -766.7, -733.3, -700.0, -666.7, -633.3, -600.0, -566.7, -533.3, -500.0, -500.0, -489.9, -479.7, -469.6, -459.5, -449.4, -439.2, -429.1, -419.0, -408.9, -398.7, -388.6, -378.5, -368.4, -358.2, -348.1, -338.0, -327.8, - |

| | | |
|---|---|---|
| | | 317.7, -307.6, -297.5, -287.3, -277.2, -267.1, -257.0, -246.8, -236.7, -226.6, -216.5, -206.3, -196.2, -186.1, -175.9, -165.8, -155.7, -145.6, -135.4, -125.3, -115.2, -105.1, -94.9, -84.8, -74.7, -64.6, -54.4, -44.3, -34.2, -24.1, -13.9, -3.8, 6.3, 16.5, 26.6, 36.7, 46.8, 57.0, 67.1, 77.2, 87.3, 97.5, 107.6, 117.7, 127.8, 138.0, 148.1, 158.2, 168.4, 178.5, 188.6, 198.7, 208.9, 219.0, 229.1, 239.2, 249.4, 259.5, 269.6, 279.7, 289.9, 300.0, 300.0, 333.3, 366.7, 400.0, 433.3, 466.7, 500.0, 533.3, 566.7, 600.0} |
| | | [30] {-800.0, -766.7, -733.3, -700.0, -666.7, -633.3, -600.0, -566.7, -533.3, -500.0, -500.0, -489.9, -479.7, -469.6, -459.5, -449.4, -439.2, -429.1, -419.0, -408.9, -398.7, -388.6, -378.5, -368.4, -358.2, -348.1, -338.0, -327.8, -317.7, -307.6, -297.5, -287.3, -277.2, -267.1, -257.0, -246.8, -236.7, -226.6, -216.5, -206.3, -196.2, -186.1, -175.9, -165.8, -155.7, -145.6, -135.4, -125.3, -115.2, -105.1, -94.9, -84.8, -74.7, -64.6, -54.4, -44.3, -34.2, -24.1, -13.9, -3.8, 6.3, 16.5, 26.6, 36.7, 46.8, 57.0, 67.1, 77.2, 87.3, 97.5, 107.6, 117.7, 127.8, 138.0, 148.1, 158.2, 168.4, 178.5, 188.6, 198.7, 208.9, 219.0, 229.1, 239.2, 249.4, 259.5, 269.6, 279.7, 289.9, 300.0, 300.0, 333.3, 366.7, 400.0, 433.3, 466.7, 500.0, 533.3, 566.7, 600.0} |
| | | [40] {-800.0, -766.7, -733.3, -700.0, -666.7, -633.3, -600.0, -566.7, -533.3, -500.0, -500.0, -489.9, -479.7, -469.6, -459.5, -449.4, -439.2, -429.1, -419.0, -408.9, -398.7, -388.6, -378.5, -368.4, -358.2, -348.1, -338.0, -327.8, -317.7, -307.6, -297.5, -287.3, -277.2, -267.1, -257.0, -246.8, -236.7, -226.6, -216.5, -206.3, -196.2, -186.1, -175.9, -165.8, -155.7, -145.6, -135.4, -125.3, -115.2, -105.1, -94.9, -84.8, -74.7, -64.6, -54.4, -44.3, -34.2, -24.1, -13.9, -3.8, 6.3, 16.5, 26.6, 36.7, 46.8, 57.0, 67.1, 77.2, 87.3, 97.5, 107.6, 117.7, 127.8, 138.0, 148.1, 158.2, 168.4, 178.5, 188.6, 198.7, 208.9, 219.0, 229.1, 239.2, 249.4, 259.5, 269.6, 279.7, 289.9, 300.0, 300.0, 333.3, 366.7, 400.0, 433.3, 466.7, 500.0, 533.3, 566.7, 600.0} |
| | | [50] {-800.0, -766.7, -733.3, -700.0, -666.7, -633.3, -600.0, -566.7, -533.3, -500.0, -500.0, -489.9, -479.7, -469.6, -459.5, -449.4, -439.2, -429.1, -419.0, -408.9, -398.7, -388.6, -378.5, -368.4, -358.2, -348.1, -338.0, -327.8, -317.7, -307.6, -297.5, -287.3, -277.2, -267.1, -257.0, -246.8, -236.7, -226.6, -216.5, -206.3, -196.2, -186.1, -175.9, -165.8, -155.7, -145.6, -135.4, -125.3, -115.2, -105.1, -94.9, -84.8, -74.7, -64.6, -54.4, -44.3, -34.2, -24.1, -13.9, -3.8, 6.3, 16.5, 26.6, 36.7, 46.8, 57.0, 67.1, 77.2, 87.3, 97.5, 107.6, 117.7, 127.8, 138.0, 148.1, 158.2, 168.4, 178.5, 188.6, 198.7, 208.9, 219.0, 229.1, 239.2, 249.4, 259.5, 269.6, 279.7, 289.9, 300.0, 300.0, 333.3, 366.7, 400.0, 433.3, 466.7, 500.0, 533.3, 566.7, 600.0} |
| TAR-C24U25A35 (-$Mg^{2+}$) | U23-C6 | [5] {-1000.0, -922.2, -844.4, -766.7, -688.9, -611.1, -533.3, -455.6, -377.8, -300.0, -300.0, -289.9, -279.7, -269.6, -259.5, -249.4, -239.2, -229.1, -219.0, -208.9, -198.7, -188.6, -178.5, -168.4, -158.2, -148.1, -138.0, -127.8, -117.7, -107.6, -97.5, -87.3, -77.2, -67.1, -57.0, -46.8, -36.7, -26.6, -16.5, -6.3, 3.8, 13.9, 24.1, 34.2, 44.3, 54.4, 64.6, 74.7, 84.8, 94.9, 105.1, 115.2, 125.3, 135.4, 145.6, 155.7, 165.8, 175.9, 186.1, 196.2, 206.3, 216.5, 226.6, 236.7, 246.8, 257.0, 267.1, 277.2, 287.3, 297.5, 307.6, 317.7, 327.8, 338.0, 348.1, 358.2, 368.4, 378.5, 388.6, 398.7, 408.9, 419.0, 429.1, 439.2, 449.4, 459.5, 469.6, 479.7, 489.9, 500.0, 500.0, 555.6, 611.1, 666.7, 722.2, 777.8, 833.3, 888.9, 944.4, 1000.0} |
| | | [10] {-1000.0, -922.2, -844.4, -766.7, -688.9, -611.1, -533.3, -455.6, -377.8, -300.0, -300.0, -289.9, -279.7, -269.6, -259.5, -249.4, -239.2, -229.1, -219.0, -208.9, -198.7, -188.6, -178.5, -168.4, -158.2, -148.1, -138.0, -127.8, -117.7, -107.6, -97.5, -87.3, -77.2, -67.1, -57.0, -46.8, -36.7, -26.6, -16.5, -6.3, 3.8, 13.9, 24.1, 34.2, 44.3, 54.4, 64.6, 74.7, 84.8, 94.9, 105.1, 115.2, 125.3, 135.4, 145.6, 155.7, 165.8, 175.9, 186.1, 196.2, 206.3, 216.5, 226.6, 236.7, 246.8, 257.0, 267.1, 277.2, 287.3, 297.5, 307.6, 317.7, 327.8, 338.0, 348.1, 358.2, 368.4, 378.5, 388.6, 398.7, 408.9, 419.0, 429.1, 439.2, 449.4, 459.5, 469.6, 479.7, 489.9, 500.0, 500.0, 555.6, 611.1, 666.7, 722.2, 777.8, 833.3, 888.9, 944.4, 1000.0} |

| | | |
|---|---|---|
| | | [25] {-1000.0, -922.2, -844.4, -766.7, -688.9, -611.1, -533.3, -455.6, -377.8, -300.0, -300.0, -289.9, -279.7, -269.6, -259.5, -249.4, -239.2, -229.1, -219.0, -208.9, -198.7, -188.6, -178.5, -168.4, -158.2, -148.1, -138.0, -127.8, -117.7, -107.6, -97.5, -87.3, -77.2, -67.1, -57.0, -46.8, -36.7, -26.6, -16.5, -6.3, 3.8, 13.9, 24.1, 34.2, 44.3, 54.4, 64.6, 74.7, 84.8, 94.9, 105.1, 115.2, 125.3, 135.4, 145.6, 155.7, 165.8, 175.9, 186.1, 196.2, 206.3, 216.5, 226.6, 236.7, 246.8, 257.0, 267.1, 277.2, 287.3, 297.5, 307.6, 317.7, 327.8, 338.0, 348.1, 358.2, 368.4, 378.5, 388.6, 398.7, 408.9, 419.0, 429.1, 439.2, 449.4, 459.5, 469.6, 479.7, 489.9, 500.0, 500.0, 555.6, 611.1, 666.7, 722.2, 777.8, 833.3, 888.9, 944.4, 1000.0} |
| | | [50] {-1000.0, -922.2, -844.4, -766.7, -688.9, -611.1, -533.3, -455.6, -377.8, -300.0, -300.0, -289.9, -279.7, -269.6, -259.5, -249.4, -239.2, -229.1, -219.0, -208.9, -198.7, -188.6, -178.5, -168.4, -158.2, -148.1, -138.0, -127.8, -117.7, -107.6, -97.5, -87.3, -77.2, -67.1, -57.0, -46.8, -36.7, -26.6, -16.5, -6.3, 3.8, 13.9, 24.1, 34.2, 44.3, 54.4, 64.6, 74.7, 84.8, 94.9, 105.1, 115.2, 125.3, 135.4, 145.6, 155.7, 165.8, 175.9, 186.1, 196.2, 206.3, 216.5, 226.6, 236.7, 246.8, 257.0, 267.1, 277.2, 287.3, 297.5, 307.6, 317.7, 327.8, 338.0, 348.1, 358.2, 368.4, 378.5, 388.6, 398.7, 408.9, 419.0, 429.1, 439.2, 449.4, 459.5, 469.6, 479.7, 489.9, 500.0, 500.0, 555.6, 611.1, 666.7, 722.2, 777.8, 833.3, 888.9, 944.4, 1000.0} |
| TAR-C24U25A35 (-$Mg^{2+}$) | U38-N3 | [10] {-1000.0, -944.4, -888.9, -833.3, -777.8, -722.2, -666.7, -611.1, -555.6, -500.0, -500.0, -489.9, -479.7, -469.6, -459.5, -449.4, -439.2, -429.1, -419.0, -408.9, -398.7, -388.6, -378.5, -368.4, -358.2, -348.1, -338.0, -327.8, -317.7, -307.6, -297.5, -287.3, -277.2, -267.1, -257.0, -246.8, -236.7, -226.6, -216.5, -206.3, -196.2, -186.1, -175.9, -165.8, -155.7, -145.6, -135.4, -125.3, -115.2, -105.1, -94.9, -84.8, -74.7, -64.6, -54.4, -44.3, -34.2, -24.1, -13.9, -3.8, 6.3, 16.5, 26.6, 36.7, 46.8, 57.0, 67.1, 77.2, 87.3, 97.5, 107.6, 117.7, 127.8, 138.0, 148.1, 158.2, 168.4, 178.5, 188.6, 198.7, 208.9, 219.0, 229.1, 239.2, 249.4, 259.5, 269.6, 279.7, 289.9, 300.0, 300.0, 377.8, 455.6, 533.3, 611.1, 688.9, 766.7, 844.4, 922.2, 1000.0} |
| | | [20] {-1000.0, -944.4, -888.9, -833.3, -777.8, -722.2, -666.7, -611.1, -555.6, -500.0, -500.0, -489.9, -479.7, -469.6, -459.5, -449.4, -439.2, -429.1, -419.0, -408.9, -398.7, -388.6, -378.5, -368.4, -358.2, -348.1, -338.0, -327.8, -317.7, -307.6, -297.5, -287.3, -277.2, -267.1, -257.0, -246.8, -236.7, -226.6, -216.5, -206.3, -196.2, -186.1, -175.9, -165.8, -155.7, -145.6, -135.4, -125.3, -115.2, -105.1, -94.9, -84.8, -74.7, -64.6, -54.4, -44.3, -34.2, -24.1, -13.9, -3.8, 6.3, 16.5, 26.6, 36.7, 46.8, 57.0, 67.1, 77.2, 87.3, 97.5, 107.6, 117.7, 127.8, 138.0, 148.1, 158.2, 168.4, 178.5, 188.6, 198.7, 208.9, 219.0, 229.1, 239.2, 249.4, 259.5, 269.6, 279.7, 289.9, 300.0, 300.0, 377.8, 455.6, 533.3, 611.1, 688.9, 766.7, 844.4, 922.2, 1000.0} |
| | | [25] {-1000.0, -944.4, -888.9, -833.3, -777.8, -722.2, -666.7, -611.1, -555.6, -500.0, -500.0, -489.9, -479.7, -469.6, -459.5, -449.4, -439.2, -429.1, -419.0, -408.9, -398.7, -388.6, -378.5, -368.4, -358.2, -348.1, -338.0, -327.8, -317.7, -307.6, -297.5, -287.3, -277.2, -267.1, -257.0, -246.8, -236.7, -226.6, -216.5, -206.3, -196.2, -186.1, -175.9, -165.8, -155.7, -145.6, -135.4, -125.3, -115.2, -105.1, -94.9, -84.8, -74.7, -64.6, -54.4, -44.3, -34.2, -24.1, -13.9, -3.8, 6.3, 16.5, 26.6, 36.7, 46.8, 57.0, 67.1, 77.2, 87.3, 97.5, 107.6, 117.7, 127.8, 138.0, 148.1, 158.2, 168.4, 178.5, 188.6, 198.7, 208.9, 219.0, 229.1, 239.2, 249.4, 259.5, 269.6, 279.7, 289.9, 300.0, 300.0, 377.8, 455.6, 533.3, 611.1, 688.9, 766.7, 844.4, 922.2, 1000.0} |
| | | [30] {-1000.0, -944.4, -888.9, -833.3, -777.8, -722.2, -666.7, -611.1, -555.6, -500.0, -500.0, -489.9, -479.7, -469.6, -459.5, -449.4, -439.2, -429.1, -419.0, -408.9, -398.7, -388.6, -378.5, -368.4, -358.2, -348.1, -338.0, -327.8, -317.7, -307.6, -297.5, -287.3, -277.2, -267.1, -257.0, -246.8, -236.7, -226.6, -216.5, -206.3, -196.2, -186.1, -175.9, -165.8, -155.7, -145.6, -135.4, -125.3, -115.2, -105.1, -94.9, -84.8, -74.7, -64.6, -54.4, -44.3, -34.2, -24.1, -13.9, -3.8, 6.3, 16.5, 26.6, 36.7, 46.8, 57.0, 67.1, 77.2, 87.3, 97.5, 107.6, 117.7, 127.8, 138.0, 148.1, |

| | | |
|---|---|---|
| | | 158.2, 168.4, 178.5, 188.6, 198.7, 208.9, 219.0, 229.1, 239.2, 249.4, 259.5, 269.6, 279.7, 289.9, 300.0, 300.0, 377.8, 455.6, 533.3, 611.1, 688.9, 766.7, 844.4, 922.2, 1000.0} |
| TAR-C24U25A35 (+$Mg^{2+}$) | U23-C6 | [10] {-1000.0, -922.2, -844.4, -766.7, -688.9, -611.1, -533.3, -455.6, -377.8, -300.0, -300.0, -289.9, -279.7, -269.6, -259.5, -249.4, -239.2, -229.1, -219.0, -208.9, -198.7, -188.6, -178.5, -168.4, -158.2, -148.1, -138.0, -127.8, -117.7, -107.6, -97.5, -87.3, -77.2, -67.1, -57.0, -46.8, -36.7, -26.6, -16.5, -6.3, 3.8, 13.9, 24.1, 34.2, 44.3, 54.4, 64.6, 74.7, 84.8, 94.9, 105.1, 115.2, 125.3, 135.4, 145.6, 155.7, 165.8, 175.9, 186.1, 196.2, 206.3, 216.5, 226.6, 236.7, 246.8, 257.0, 267.1, 277.2, 287.3, 297.5, 307.6, 317.7, 327.8, 338.0, 348.1, 358.2, 368.4, 378.5, 388.6, 398.7, 408.9, 419.0, 429.1, 439.2, 449.4, 459.5, 469.6, 479.7, 489.9, 500.0, 500.0, 555.6, 611.1, 666.7, 722.2, 777.8, 833.3, 888.9, 944.4, 1000.0} |
| | | [20] {-1000.0, -922.2, -844.4, -766.7, -688.9, -611.1, -533.3, -455.6, -377.8, -300.0, -300.0, -289.9, -279.7, -269.6, -259.5, -249.4, -239.2, -229.1, -219.0, -208.9, -198.7, -188.6, -178.5, -168.4, -158.2, -148.1, -138.0, -127.8, -117.7, -107.6, -97.5, -87.3, -77.2, -67.1, -57.0, -46.8, -36.7, -26.6, -16.5, -6.3, 3.8, 13.9, 24.1, 34.2, 44.3, 54.4, 64.6, 74.7, 84.8, 94.9, 105.1, 115.2, 125.3, 135.4, 145.6, 155.7, 165.8, 175.9, 186.1, 196.2, 206.3, 216.5, 226.6, 236.7, 246.8, 257.0, 267.1, 277.2, 287.3, 297.5, 307.6, 317.7, 327.8, 338.0, 348.1, 358.2, 368.4, 378.5, 388.6, 398.7, 408.9, 419.0, 429.1, 439.2, 449.4, 459.5, 469.6, 479.7, 489.9, 500.0, 500.0, 555.6, 611.1, 666.7, 722.2, 777.8, 833.3, 888.9, 944.4, 1000.0} |
| | | [30] {-1000.0, -922.2, -844.4, -766.7, -688.9, -611.1, -533.3, -455.6, -377.8, -300.0, -300.0, -289.9, -279.7, -269.6, -259.5, -249.4, -239.2, -229.1, -219.0, -208.9, -198.7, -188.6, -178.5, -168.4, -158.2, -148.1, -138.0, -127.8, -117.7, -107.6, -97.5, -87.3, -77.2, -67.1, -57.0, -46.8, -36.7, -26.6, -16.5, -6.3, 3.8, 13.9, 24.1, 34.2, 44.3, 54.4, 64.6, 74.7, 84.8, 94.9, 105.1, 115.2, 125.3, 135.4, 145.6, 155.7, 165.8, 175.9, 186.1, 196.2, 206.3, 216.5, 226.6, 236.7, 246.8, 257.0, 267.1, 277.2, 287.3, 297.5, 307.6, 317.7, 327.8, 338.0, 348.1, 358.2, 368.4, 378.5, 388.6, 398.7, 408.9, 419.0, 429.1, 439.2, 449.4, 459.5, 469.6, 479.7, 489.9, 500.0, 500.0, 555.6, 611.1, 666.7, 722.2, 777.8, 833.3, 888.9, 944.4, 1000.0} |
| | | [40] {-1000.0, -922.2, -844.4, -766.7, -688.9, -611.1, -533.3, -455.6, -377.8, -300.0, -300.0, -289.9, -279.7, -269.6, -259.5, -249.4, -239.2, -229.1, -219.0, -208.9, -198.7, -188.6, -178.5, -168.4, -158.2, -148.1, -138.0, -127.8, -117.7, -107.6, -97.5, -87.3, -77.2, -67.1, -57.0, -46.8, -36.7, -26.6, -16.5, -6.3, 3.8, 13.9, 24.1, 34.2, 44.3, 54.4, 64.6, 74.7, 84.8, 94.9, 105.1, 115.2, 125.3, 135.4, 145.6, 155.7, 165.8, 175.9, 186.1, 196.2, 206.3, 216.5, 226.6, 236.7, 246.8, 257.0, 267.1, 277.2, 287.3, 297.5, 307.6, 317.7, 327.8, 338.0, 348.1, 358.2, 368.4, 378.5, 388.6, 398.7, 408.9, 419.0, 429.1, 439.2, 449.4, 459.5, 469.6, 479.7, 489.9, 500.0, 500.0, 555.6, 611.1, 666.7, 722.2, 777.8, 833.3, 888.9, 944.4, 1000.0} |
| TAR-C24U25A35 (+$Mg^{2+}$) | U38-N3 | [10] {-1000.0, -944.4, -888.9, -833.3, -777.8, -722.2, -666.7, -611.1, -555.6, -500.0, -500.0, -489.9, -479.7, -469.6, -459.5, -449.4, -439.2, -429.1, -419.0, -408.9, -398.7, -388.6, -378.5, -368.4, -358.2, -348.1, -338.0, -327.8, -317.7, -307.6, -297.5, -287.3, -277.2, -267.1, -257.0, -246.8, -236.7, -226.6, -216.5, -206.3, -196.2, -186.1, -175.9, -165.8, -155.7, -145.6, -135.4, -125.3, -115.2, -105.1, -94.9, -84.8, -74.7, -64.6, -54.4, -44.3, -34.2, -24.1, -13.9, -3.8, 6.3, 16.5, 26.6, 36.7, 46.8, 57.0, 67.1, 77.2, 87.3, 97.5, 107.6, 117.7, 127.8, 138.0, 148.1, 158.2, 168.4, 178.5, 188.6, 198.7, 208.9, 219.0, 229.1, 239.2, 249.4, 259.5, 269.6, 279.7, 289.9, 300.0, 300.0, 377.8, 455.6, 533.3, 611.1, 688.9, 766.7, 844.4, 922.2, 1000.0} |
| | | [20] {-1000.0, -944.4, -888.9, -833.3, -777.8, -722.2, -666.7, -611.1, -555.6, -500.0, -500.0, -489.9, -479.7, -469.6, -459.5, -449.4, -439.2, -429.1, -419.0, -408.9, -398.7, -388.6, -378.5, -368.4, -358.2, -348.1, -338.0, -327.8, -317.7, -307.6, -297.5, -287.3, -277.2, -267.1, -257.0, -246.8, -236.7, -226.6, -216.5, -206.3, -196.2, -186.1, -175.9, -165.8, -155.7, -145.6, -135.4, -125.3, -115.2, -105.1, -94.9, -84.8, -74.7, -64.6, -54.4, -44.3, -34.2, |

| | | |
|---|---|---|
| | | -24.1, -13.9, -3.8, 6.3, 16.5, 26.6, 36.7, 46.8, 57.0, 67.1, 77.2, 87.3, 97.5, 107.6, 117.7, 127.8, 138.0, 148.1, 158.2, 168.4, 178.5, 188.6, 198.7, 208.9, 219.0, 229.1, 239.2, 249.4, 259.5, 269.6, 279.7, 289.9, 300.0, 300.0, 377.8, 455.6, 533.3, 611.1, 688.9, 766.7, 844.4, 922.2, 1000.0} |
| | | [30] {-1000.0, -944.4, -888.9, -833.3, -777.8, -722.2, -666.7, -611.1, -555.6, -500.0, -500.0, -489.9, -479.7, -469.6, -459.5, -449.4, -439.2, -429.1, -419.0, -408.9, -398.7, -388.6, -378.5, -368.4, -358.2, -348.1, -338.0, -327.8, -317.7, -307.6, -297.5, -287.3, -277.2, -267.1, -257.0, -246.8, -236.7, -226.6, -216.5, -206.3, -196.2, -186.1, -175.9, -165.8, -155.7, -145.6, -135.4, -125.3, -115.2, -105.1, -94.9, -84.8, -74.7, -64.6, -54.4, -44.3, -34.2, -24.1, -13.9, -3.8, 6.3, 16.5, 26.6, 36.7, 46.8, 57.0, 67.1, 77.2, 87.3, 97.5, 107.6, 117.7, 127.8, 138.0, 148.1, 158.2, 168.4, 178.5, 188.6, 198.7, 208.9, 219.0, 229.1, 239.2, 249.4, 259.5, 269.6, 279.7, 289.9, 300.0, 300.0, 377.8, 455.6, 533.3, 611.1, 688.9, 766.7, 844.4, 922.2, 1000.0} |
| | | [40] {-1000.0, -944.4, -888.9, -833.3, -777.8, -722.2, -666.7, -611.1, -555.6, -500.0, -500.0, -489.9, -479.7, -469.6, -459.5, -449.4, -439.2, -429.1, -419.0, -408.9, -398.7, -388.6, -378.5, -368.4, -358.2, -348.1, -338.0, -327.8, -317.7, -307.6, -297.5, -287.3, -277.2, -267.1, -257.0, -246.8, -236.7, -226.6, -216.5, -206.3, -196.2, -186.1, -175.9, -165.8, -155.7, -145.6, -135.4, -125.3, -115.2, -105.1, -94.9, -84.8, -74.7, -64.6, -54.4, -44.3, -34.2, -24.1, -13.9, -3.8, 6.3, 16.5, 26.6, 36.7, 46.8, 57.0, 67.1, 77.2, 87.3, 97.5, 107.6, 117.7, 127.8, 138.0, 148.1, 158.2, 168.4, 178.5, 188.6, 198.7, 208.9, 219.0, 229.1, 239.2, 249.4, 259.5, 269.6, 279.7, 289.9, 300.0, 300.0, 377.8, 455.6, 533.3, 611.1, 688.9, 766.7, 844.4, 922.2, 1000.0} |
